# Faster but not sweeter: A model of *E. coli* Re-level lipopolysaccharide for Martini 3 and a Martini 2 version with accelerated kinetics

**DOI:** 10.1101/2024.05.27.596027

**Authors:** Astrid F. Brandner, Dheeraj Prakaash, Alexandre Blanco González, Fergus Waterhouse, Syma Khalid

## Abstract

Lipopolysaccharide (LPS) is a complex glycolipid molecule that is the main lipidic component of the outer leaflet of the outer membrane of Gram-negative bacteria. It has very limited lateral motion compared to phospholipids, which are more ubiquitous in biological membranes, including in the inner leaflet of the outer membrane of Gram-negative bacteria. The slow-moving nature of LPS can present a hurdle for molecular dynamics simulations given the (pragmatically) accessible timescales to simulations are currently limited to microseconds, during which LPS displays some conformational dynamics, but hardly any lateral diffusion. Thus, it is not feasible to observe phenomena such as insertion of molecules, including antibiotics/antimicrobials directly into the outer membrane from the extracellular side nor to observe LPS dissociating from proteins *via* molecular dynamics using currently available models at the atomistic and more coarse-grained levels of granularity. Here we present a model of deep rough LPS compatible with the Martini 2 coarse-grained force field with scaled down non-bonded interactions to enable faster diffusion. We show that the faster-diffusing LPS model is able to reproduce the salient biophysical properties of the standard models, but due to its faster lateral motion, molecules are able to penetrate deeper into membranes containing the faster model. We show that faster LPS model is able to reproduce experimentally determined patterns of interaction with outer membrane proteins, while also allowing for LPS to associate/dissociate with proteins within microsecond timescales. We also complete the Martini 3 LPS toolkit for *E. coli* by presenting a (standard) model of deep rough LPS for this forcefield.

## 1 INTRODUCTION

Lipopolysaccharide (LPS) is a major component of the outer membranes of Gram-negative bacteria, and indeed the outer leaflet of outer membranes contain LPS almost exclusively as the lipidic component ^1^. It is also found in marine-derived aerosols ^2–4^. The complex biochemistry of LPS furnishes it with the ability to form multiple types of non-covalent interactions with other molecules as well as itself. In summary, the acyl tails (there are usually 4 to 6 depending on the bacterial species) can form hydrophobic interactions with similar chemical moieties, whereas the sugars can form multiple hydrogen bonds with polar molecules, and those sugars that are phosphorylated, have additional electrostatic interaction capabilities. The multiple interactions neighbouring LPS molecules form with each other, the cross-linking of the phosphorylated regions by divalent cations, as well as the large mass of the molecule, all contribute to the limited lateral mobility of LPS ^5^. Indeed, the limited diffusion of LPS is a major contributor to the impermeability of bacterial outer membranes and the difficulty of penetrating these membranes with antimicrobials ^6–8^. Before discussing simulations of LPS, it is useful to describe the molecule in more detail. Full length LPS is composed of lipid A whose acyl chains face the tails of the inner leaflet phospholipids. Lipid A consists of acyl chains attached to a glucosamine disaccharide, each of the sugars usually contains a single phosphate group. Covalently linked to lipid A are core oligosaccharides, whichSx in turn are connected to highly diverse O-antigen polysaccharides, the latter can vary in their number of repeating units (up to ∼100) ^9^. It is important to note that different strains of even the same bacterium can vary in the length of their LPS. ReLPS (also referred to as deep rough) corresponds to lipid A + two Kdo sugars, and is the minimal viable form of LPS in most Gram-negative bacteria. RaLPS (also referred to as rough LPS) is the name given to the length of LPS that contains all of the core oligosaccharides, but not the O-antigen region ^10^. Molecular dynamics (MD) simulation studies of LPS can be broadly divided into two categories; those that consider single LPS molecules and those in which LPS-containing bilayers are simulated. The former category includes studies of LPS recognition by host elements such as *via* the Toll-like receptor (TLR-4) in humans ^11, 12^. In such cases, the interaction between a single LPS molecule and the receptor is studied with the simulation initiated with the LPS already bound to the receptor, thus the encounter between the two molecules is not considered and the slow diffusion of LPS is not a problem. However, it becomes more problematic when multiple LPS molecules are interacting with each other (and other molecules) within a lipid bilayer. Here, the slow lateral motion of LPS necessitates extended equilibration periods compared to simpler phospholipid-containing bilayers. Furthermore, the kinetic barriers to permeation of molecules such as antimicrobial peptides into the LPS-containing bilayers are unsurmountable within the microsecond timescales accessible to MD simulations, despite these peptides being known experimentally to partition into the bilayers ^13^. Coarse-grained (CG) models offer a speed up compared to all-atom or united-atom models, however even with the most popular near-atomistic resolution (i.e. not mesoscale) coarse-grained force-fields (e.g. Martini ^14, 15^), breaking beyond the microsecond timescale currently remains impractical. While algorithms for enhanced sampling such as metadynamics and umbrella sampling do offer routes to accessing slower events, we have previously shown that for simulations involving LPS-containing bilayers this is not straightforward ^5^.

Consequently, there is a need for LPS models that exhibit accelerated lateral motion. To advance the currently available coarse-grained toolkit for modelling LPS, here we present two models of ReLPS. Firstly, a standard model of ReLPS that is compatible with the Martini 3 forcefield, given this was omitted from a recently reported Martini 3 parameter set for other levels of LPS ^16^, and secondly an accelerated or ‘fast’ ReLPS model that is compatible with the popular Martini 2 coarse-grained suite. Models for both force-fields are required, given Martini 2 has the larger repertoire of molecules including lipids and sugars available, whereas Martini 3 is the latest parameter set, however the range of systems that can be simulated is more limited due to missing parameters for some key molecules (e.g. many lipids and disaccharides). The fast ReLPS model has ∼ 29 times faster lateral diffusion compared to the standard Martini 2 ReLPS. We show that the faster diffusion allows deeper penetration of the antimicrobial polymyxin B1 without rendering the bilayer overly penetrable. Furthermore, we show that our ‘fast’ ReLPS model recapitulates known outer membrane protein-LPS interactions starting from unbiased system configurations, crucially also allowing for protein-LPS dissociation.

## 2 METHODS

Atomistic simulations of symmetric ReLPS bilayers were performed with the CHARMM36m force field^17^ to parametrise the new Martini 3 ReLPS. The coarse-grained molecular dynamics (CGMD) simulations presented in this work were performed with the Martini force field version 2.2 ^14^ or 3.0 ^18^. Protein structures used in CGMD simulations were coarse-grained using the martinize.py or martinize2 script for the Martini 2.2 or Martini 3.0 force fields respectively. An *ad hoc* version of the insane.py script ^19^ was used to insert proteins into model membranes. The modifications to the insane.py script include the original ReLPS parameters according to the Martini 2.2 force field as provided by CHARMM-GUI Martini Maker ^15^, and our newly parameterised ReLPS (Martini 3) and fast ReLPS (Martini 2). See **table 1** for details on all the simulations presented in this work.

**Table 1.**
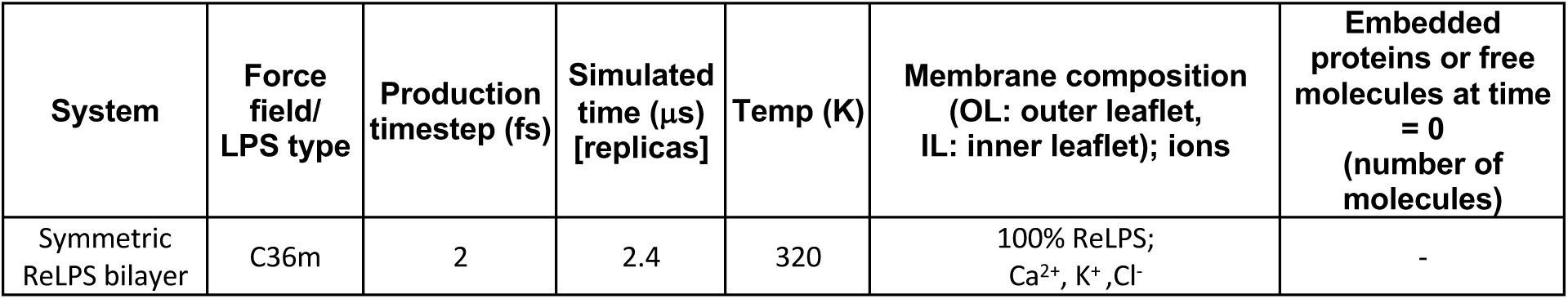
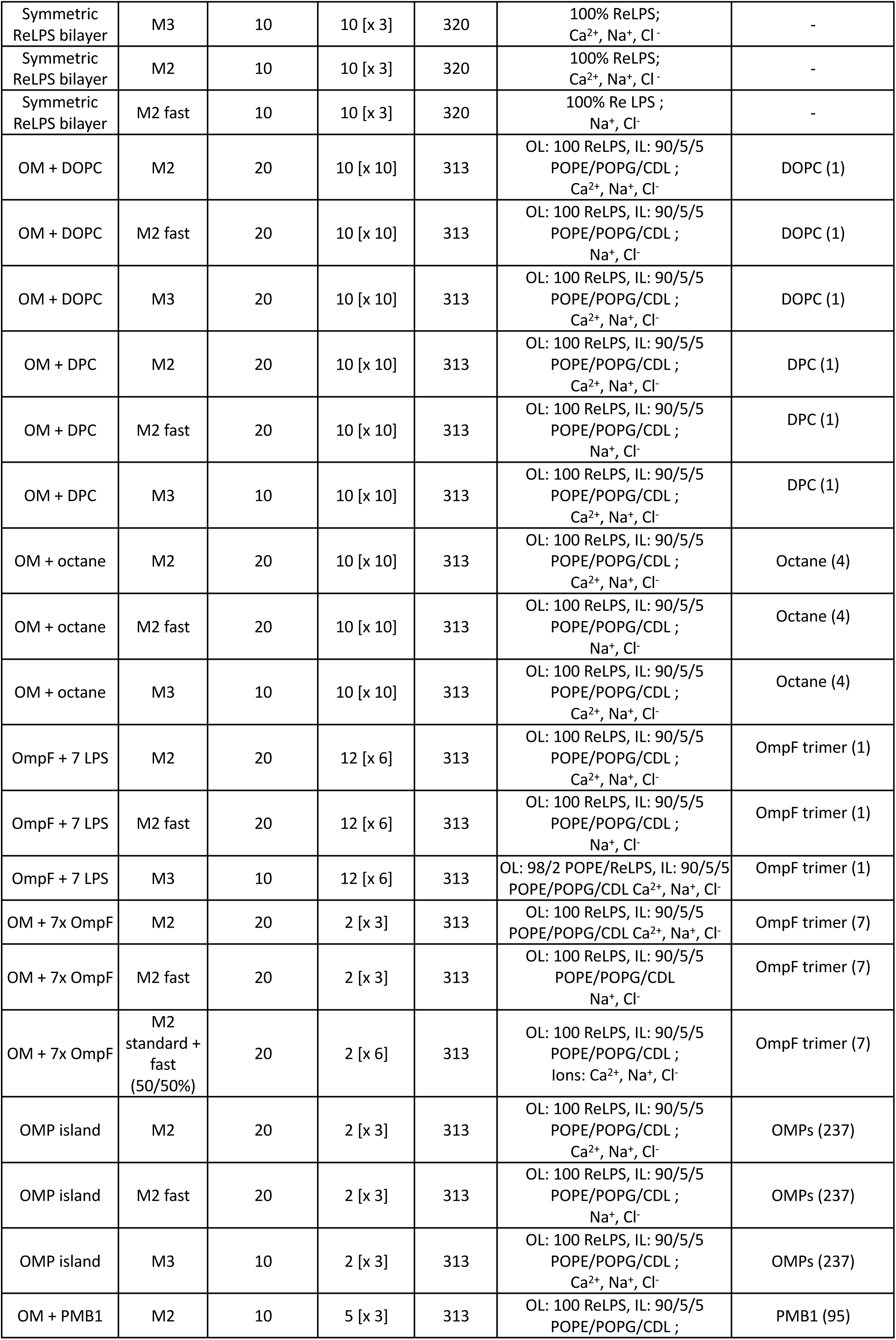
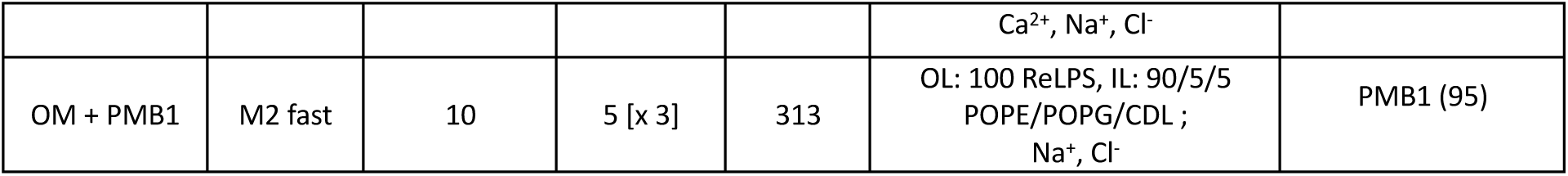
Details on simulation setup for all systems conducted in this work. . The abbreviations are as follows: DPC: dodecylphosphorylcholine, POPE: palmitoyl-2-oleoyl-sn-glycero-3-phosphoethanolamine, POPG: 1-palmitoyl-2-oleoyl-sn-glycero-3-phosphoglycerol, CDL: cardiolipin, PMB1: polymyxin B1, OM: outer membrane, OMP: outer membrane protein, OL: outer leaflet, IL: inner leaflet. Force-field abbreviations are: C36m: CHARMM36m, M2: Martini 2 standard, M2 fast: fast Martini 2, M3: Martini 3.

### 2.1 Atomistic simulations and Martini 3 parameterisation

Atomistic simulations of a symmetric ReLPS bilayer were set up with CHARMM-GUI ^15^ using the CHARMM36m forcefield. The initial box size was 15x15x19 nm^3^ containing the membrane (119 ReLPS molecules in each leaflet) solvated in 0.15 M KCl solution. 714 Ca^2+^ ions were included in the ReLPS headgroup and core regions to neutralise charges. The standard protocol of CHARMM-GUI was followed for minimization and equilibration in the NPT ensemble (320 K, 1 bar). A detailed description can be found in the supplementary methods. The production run was conducted for 2.4 μs.

Coarse-grained coordinates and initial bonded parameters were generated from the atomistic simulation following the mapping scheme showed in **fig 1** *via* PyCGTOOL ^20^. Briefly, first a mapping scheme from all-atom to coarse-grain was chosen to define the CG topology and to map coordinates from the reference atomistic trajectory. Secondly, coarse-grained bonded parameters (equilibrium distances and force constants) were obtained *via* the modified Boltzmann Inversion technique implemented in PyCGTOOL. A detailed atom list from all-atom to CG mapping and the final parameters are provided in **tables S1-3** and additional text within the **supplementary material**. A time step of 10 fs was chosen a consequence of the resolution of the mapping scheme. In the current version, certain “tiny beads” (i.e. those beads with mass = 36 a.m.u, that represent 2 heavy atoms in Martini 3) are connected via a bond potential and not a constraint, producing a bond with an estimated oscillation period of e.g 0.15 ps which is smaller than ten times the timestep, which should be avoided for energy conservation between time steps. We note here that according to Berendsen and co-workers, a general guideline for the Verlet leap-frog integration scheme is a minimum of five numerical integration steps per period of harmonic oscillation for integration with reasonable accuracy ^21^.

**Figure 1.**
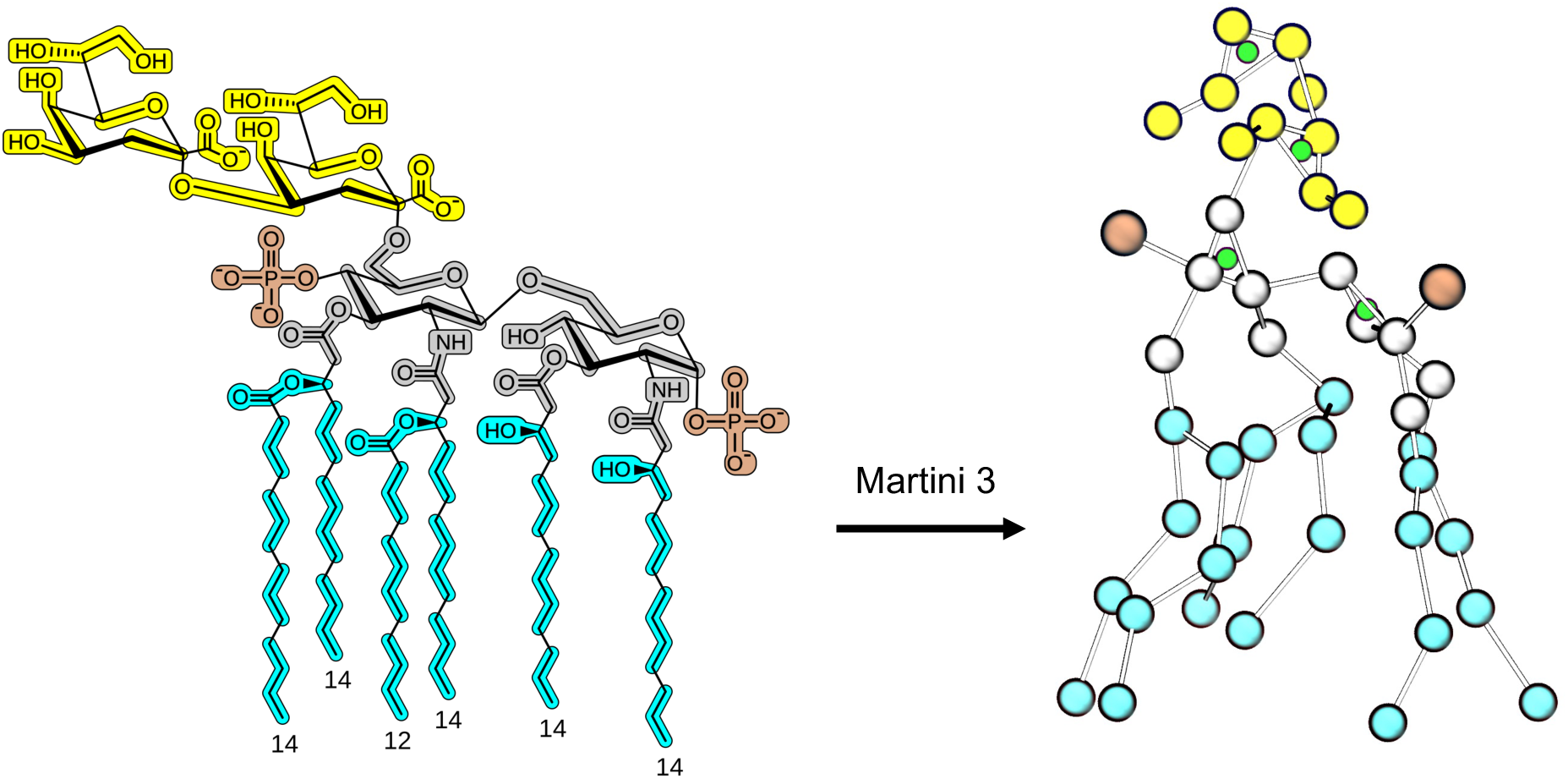
Martini 3 ReLPS mapping. Groups of two to four heavy atoms were mapped into coarse-grained beads. Notably the sugar rings were modelled with an extra virtual site bead (depicted in green) to better characterise their volume and interactions. Here, each highlighted segment of the 2D depiction corresponds to the same-coloured bead on the right CG molecule taken from a simulation snapshot. The four different colors represent: Ketodeoxyoctonic acid (Kdo) sugars i.e. first two in core region (yellow), D-Glucosamine (GlcN) sugars from lipid A moiety (grey), phosphate substituents (brown), acyl chains (cyan). A detailed atom list from all-atom to CG mapping is given in **table S1**, and the bead types annotated in **fig S1**.

Martini 3 parameters for dodecyl phosphocholine (DPC) implemented here are according to those proposed by Cabezudo *et al*. ^22^.

### 2.2 Coarse-grained simulations

#### 2.2.1 General simulation protocols

All CGMD simulation systems were solvated with the (standard) Martini water model before adding 0.15 M NaCl. Charges were neutralised by replacing water beads with either divalent or monovalent cations or both according to the concentration (0/50/100%) of ReLPS and the type of ReLPS (standard/ fast Martini 2) used in the membrane. All simulation systems with 100% standard (Martini 2 or 3) ReLPS in the outer leaflet were neutralised with divalent cations (Ca^2+^) ions in addition to 0.15 M NaCl, whereas 100% fast (Martini 2) ReLPS-containing systems were neutralised with monovalent cations (Na^+^) in addition to 0.15 M NaCl.

##### Martini 2

Details of energy minimisation and equilibration are provided in the supplementary methods. For production simulations, at least 3 replicas were performed each initiated from the end of the equilibration stage. The simulation protocols for equilibration, details of which are given in the supplementary methods section were maintained for the production simulations, but for Martini 2 simulations the pressure coupling was achieved *via* the Parrinello-Rahman barostat ^23^ with a time constant of 12 ps.

##### Martini 3

Simulations performed with the Martini 3 force field were equilibrated after minimisation as stated in the supplementary methods section. In contrast with simulations with Martini 2, the reaction-field method was used to treat electrostatic interactions. At least 3 replicas were performed for production simulations with a 10 fs timestep using the Parrinello-Rahman barostat. They were initiated from the end of the equilibration stage.

#### 2.2.2 *E. coli* outer membrane model composition

The composition of the outer membrane in CGMD simulations reported here was as follows: Palmitoyl-2-oleoyl-sn-glycero-3-phosphoethanolamine (POPE) / 1-palmitoyl-2-oleoyl-sn-glycero-3-phosphoglycerol (POPG) / cardiolipin with a −2 net charge (CDL2): 90/5/5 for the inner leaflet and 100% ReLPS in the outer leaflet. A special set of CGMD simulations contained 2% ReLPS and 98% POPE in the outer leaflet to help accelerate diffusion of lipids to better assess binding and unbinding of ReLPS with outer membrane proteins (OMPs). The X-ray crystal structure of OmpF used in this work was resolved at 1.98 Å (PDB: 3K1B) ^24^.

#### 2.2.3 Membrane permeability tests with octane, DPC and DOPC

Simulations were performed to test the spontaneous insertion of either octane, dodecyl phosphorylcholine (DPC), or dioleoyl-sn-glycero-3-phosphocholine (DOPC) in standard and fast Martini 2 ReLPS-containing outer membranes. Octane, DPC or DOPC molecules were initially placed in the solvent phase proximal to the membrane surface, but without direct contact with the membrane. Four octane molecules were simulated in each simulation replica, whereas each simulation of DPC or DOPC insertion contained only 1 molecule to avoid potential micelle formation. Position restraints were applied to octane/ DPC/ DOPC during the equilibration stage. During production, a flat-bottomed potential of 1000 kJ/mol/nm was applied to all their beads 5 nm from their equilibrated positions along the Z-coordinate to prevent them from inserting into the inner leaflet containing phospholipids. 10 production simulations of 10 μs each were performed for octane, DOPC and DPC.

#### 2.2.4 PMB1 simulations with Martini 2

To study the effect of our fast ReLPS parameters on the interactions of polymyxin B1 (PMB1) with the outer membrane, a 20 x 20 nm^2^ membrane composed of 100% ReLPS in the outer leaflet and a mixture of 90% POPE + 5% POPG + 5% cardiolipin in the inner leaflet was constructed using the online CHARMM-GUI Martini Maker ^25^. Charges were neutralised using divalent cations and 0.15 M NaCl was added to the system. To this membrane, multiple PMB1 were added in random positions and orientations in the aqueous phase above the outer leaflet such that a ratio of 2.6 ReLPS to 1 PMB1 was obtained, following one of the proportions reported in a previous study ^8^. The charges introduced by the peptide were neutralised using Cl^−^ ions.

This system was equilibrated following the standard protocol provided by CHARMM-GUI and was adapted to the presence of PMB1 in the system. Specifically, positions restraints were added to the peptides so that they would remain in the solvent and not diffuse towards the membrane during the equilibration steps. Once the system reached a state that allowed simulations to run without numerical errors at an appropriate timestep, three starting points were generated for further simulations: The first starting point remained unchanged from the original system, while the second starting point involved the replacement of half of the standard ReLPS molecules with fast ReLPS, ensuring a uniform distribution. Lastly, the third starting point featured a membrane where all standard ReLPS molecules were substituted with fast ReLPS. As mentioned previously, it is important to note that in simulations where fast ReLPS was introduced, an equivalent amount of divalent cations had to be exchanged for monovalent cations due to differences in ReLPS parameterisation. This procedure ensured that the three simulations (3 replicas each) began from identical particle configurations, with the only difference being the presence and quantity of fast ReLPS. Subsequently, these three systems were prepared for a 5 µs simulation without any restraints applied to the system.

#### 2.2.5 Simulations with multiple OmpFs

We setup simulations containing 7 OmpF trimers arranged in a hexagonal lattice based on atomic force microscopy (AFM) experiments ^26^, but placed at approximately 13 nm from each other. They were inserted into a membrane containing 100% ReLPS concentration (with an area per ReLPS lipid initially set to 1.3 nm using the insane.py script) in the outer leaflet. In addition, we performed a set of simulations containing 50% standard and 50% fast ReLPS where one setup contained a uniform distribution of standard and fast ReLPS, and the other had a non-uniform distribution. In these 50% standard/fast ReLPS-containing simulations, a sufficient number of divalent cations had to be replaced with monovalent cations in addition to the 0.15 M NaCl salt concentration due to the requirement of our fast ReLPS model.

Details of analyses and visualisation are provided in the supplementary information. All simulations in this study were performed with the GROMACS simulation suite v2019.4 or higher ^27^. Information on specific GROMACS versions used for production simulations and box dimensions for each system are provided in **table S4**.

## 3 RESULTS

### 3.1 The ReLPS model in Martini 3

The current CGMD LPS toolkit has recently been extended by the publication of an *E. coli* LPS parameter set which is compatible with Martini 3, this includes lipid A and LPS at the Ra- and O-antigen levels ^16^, but not ReLPS, which is the minimal level of LPS with which *E. coli* cells are viable. Therefore, we first sought to address this by parameterising a Martini 3 compatible model of ReLPS. Briefly, a long atomistic simulation of a pure ReLPS membrane patch neutralised with Ca^2+^ ions and in physiological salt concentration was used to obtain the first set of coordinates and bonded parameters based on the manually assigned mapping strategy (**fig 1**). The parameters are provided in the **supplementary material** as well as the probability distribution functions of each bond, angle and dihedral present in the Martini 3 model compared to their CHARMM36m counterpart. The detailed process of parameterisation is explained in the methods section. In addition, a comparison of the solvent accessible surface area (SASA) of the Martini 3 vs CHARMM36m models is provided in **fig S5.** The SASA computed for the ReLPS molecules in the context of the symmetric bilayer shows good overall agreement with its atomistic counterpart. Despite the Martini 3 SASA distribution being slightly shifted towards larger values, the difference in SASA value between Martini 3 and the atomistic reference model is 0.75 ± 2.6 nm^2^, where the propagated error is larger than the difference, thus showing good agreement between the models.

Comparison of the physical properties of this model with our previously published Martini 2 ReLPS and the reference atomistic (CHARMM36m) versions reveals good agreement and is within the same range of agreement as the aforementioned Martini 3 LPS models. The details of these properties are discussed below in the context of comparison with the subsequently parameterised ‘fast’ Martini 2 ReLPS model (**fig 3**).

### 3.2 The ‘fast’ Martini 2 ReLPS model

Given the limited lateral diffusion of LPS which is in part due to strong electrostatic interactions mediated by cross-linking of phosphorylated regions by divalent cations, we present our fast ReLPS model compatible with the Martini 2 forcefield. To parametrise this, we ran coarse-grained (Martini 2) simulations in an outer membrane model containing 100% ReLPS in the outer leaflet and phospholipids in the inner leaflet. Initially, to yield greater lateral diffusion of Martini 2 ReLPS molecules with minimal deviation from canonical physical properties of the outer membrane model, we obtained two ReLPS models where (i) the charges of its anionic beads were scaled down and neutralising divalent cations were replaced with monovalent cations, and (ii) Lennard-Jones (LJ) potentials (inter ReLPS) were scaled down (more details provided in the supplementary methods and associated **fig S6-11**). Next, we combined these models to obtain our ‘fast’ Martini 2 ReLPS model. We summarise the modified bead names and types in **fig S11A**, and the differences in the LJ potential matrices between standard and fast Martini 2 ReLPS in **fig S11B.** This ‘fast’ Martini 2 model of ReLPS was evaluated by computing its mean square displacement (MSD, revealing a ∼29-fold increase along the membrane plane) (**fig S12A**), nearest neighbour index (NNI, a measure of lipid mixing revealing an increased value by a factor larger than 4) (**fig S12B**) and partial density profiles (**fig 12C**) which showed a good agreement with those of the standard Martini 2 model.

We also simulated outer membrane models (with 7 x OmpF trimers) containing a combination of fast and standard Martini 2 ReLPS molecules (50% each) in the outer leaflet, and assessed its effect on area per lipid (APL) (**fig S13A**), membrane thickness (**fig S13B**), and MSD (**fig 13C**). These results were compared to simulations with 100% fast or 100% standard Martini 2 ReLPS. Note that we performed these simulations starting from two distinct initial setups where the fast and standard ReLPS molecules were distributed (i) uniformly or (ii) non-uniformly (**fig S13D**). From these simulations, we observed that combining fast and standard ReLPS types starting from either the uniform or non-uniform distribution did not achieve any significant increase in lateral diffusion compared to the standard ReLPS. APL and membrane thickness analyses also revealed that mixing fast and standard ReLPS types tend to provide results comparable to that of pure standard ReLPS. Therefore, from this point onward, we only simulated membranes containing either 100% standard or fast ReLPS.

Comparison of symmetric bilayers of the standard Martini 2 and fast Martini ReLPS revealed stark differences in the lateral mobility and conformational dynamics. The fast Martini 2 ReLPS exhibits about 10-fold greater lateral diffusion compared to the standard Martini 2 model (**fig 2A**). Least squares fitting of individual ReLPS molecules from different timepoints within the trajectory revealed greater conformational deviation of the fast ReLPS molecules compared to the standard model. Visual inspection indicated that while there is greater conformational lability in all regions of fast Martini 2 ReLPS molecules compared to the standard models (**fig 2B**), the difference appeared to be greater in the Kdo sugars. Overall, individual molecules of the fast Martini 2 ReLPS model exhibited a wider RMSD distribution compared to their standard Martini counterpart and reference atomistic trajectory analysed as a pseudo-Martini 2 simulation (**fig S14A, B, C**). Remarkably, RMSD distributions analysed by picking random ReLPS molecules from the pseudo-Martini 2 simulation showed that variance within each molecule was smaller compared to either Martini 2 models. However, a larger distribution was found when comparing their means (e.g. spanning values from 0.5 to 0.9 nm). Furthermore, cation interactions with the negatively charged beads of the fast ReLPS model were weakened compared to those with the standard model (**fig S15**), thereby reducing rigidity between different LPS molecules, and aiding in their individual flexibility and lateral mobility. Thus, in summary, the goal of greater motion (both lateral and the within the conformations of individual molecules) of ReLPS is realised in the new fast Martini 2 model. Interestingly, the M3 ReLPS shows similar conformational lability to the fast M2 model.

**Figure 2.**
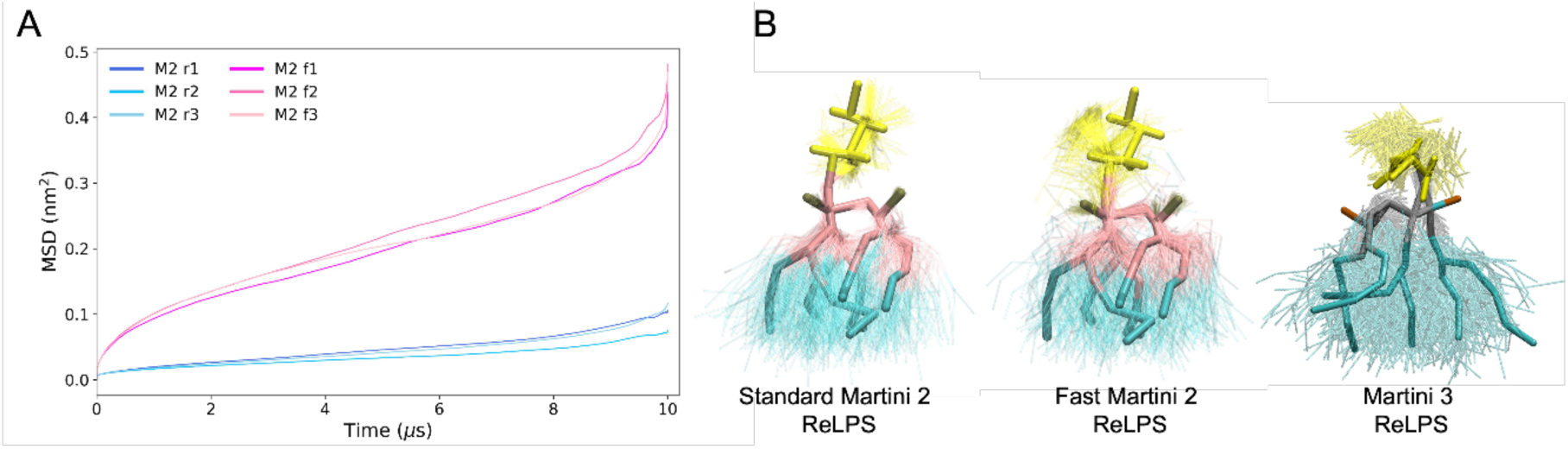
Lateral diffusion and conformational lability of the standard and fast Martini 2 ReLPS models. **(A)** A comparison between the lateral mean square displacement of individual simulation replicas of standard (M2 r1-3) and fast Martini 2 (M2 f1-3) ReLPS models. **(B)** Structural representations of ReLPS conformational flexibilities for each of the forcefields used in this work (overlaying snapshots every 100 ns after fitting in the central part of the ReLPS structure for clarity. See **fig S14A-C** for RMSD data on the structural flexibility of ReLPS and **fig S14D** for detailed MSD plots. The Martini 3 model follows the same colour scheme as in **fig 1**.

The MSDs for M3 are discussed later in this study (and plots are provided in **fig S22**) when we compare systems containing proteins - these systems are the same size for all three ReLPS models and thus provide a better comparison of the MSD of all three models.

As a thorough comparison, we quantified the biophysical properties of symmetric ReLPS bilayers using atomistic (CHARMM36m) and coarse-grained (Martini 2, fast Martini 2 and Martini 3) force fields. The membrane thickness, APL and lipid order parameters (**fig 3**) and partial density profiles (**fig S16**) of the fast Martini 2 ReLPS showed good agreement with the standard Martini 2 and atomistic models. Both Martini 2 models give values of within 4-5% of the values obtained from the atomistic simulation but mapped into Martini 2 for both thickness and APL). The Martini 3 model showed greater deviation from atomistic models (with 9% of the APL and 11% of the thickness when the atomistic trajectories are into Martini 3). Interestingly, we note water penetration into the all atom ReLPS bilayer matches more closely to the Martini 3 ReLPS bilayer density than it does to either of the two Martini 2 datasets (**fig S16**).

**Figure 3.**
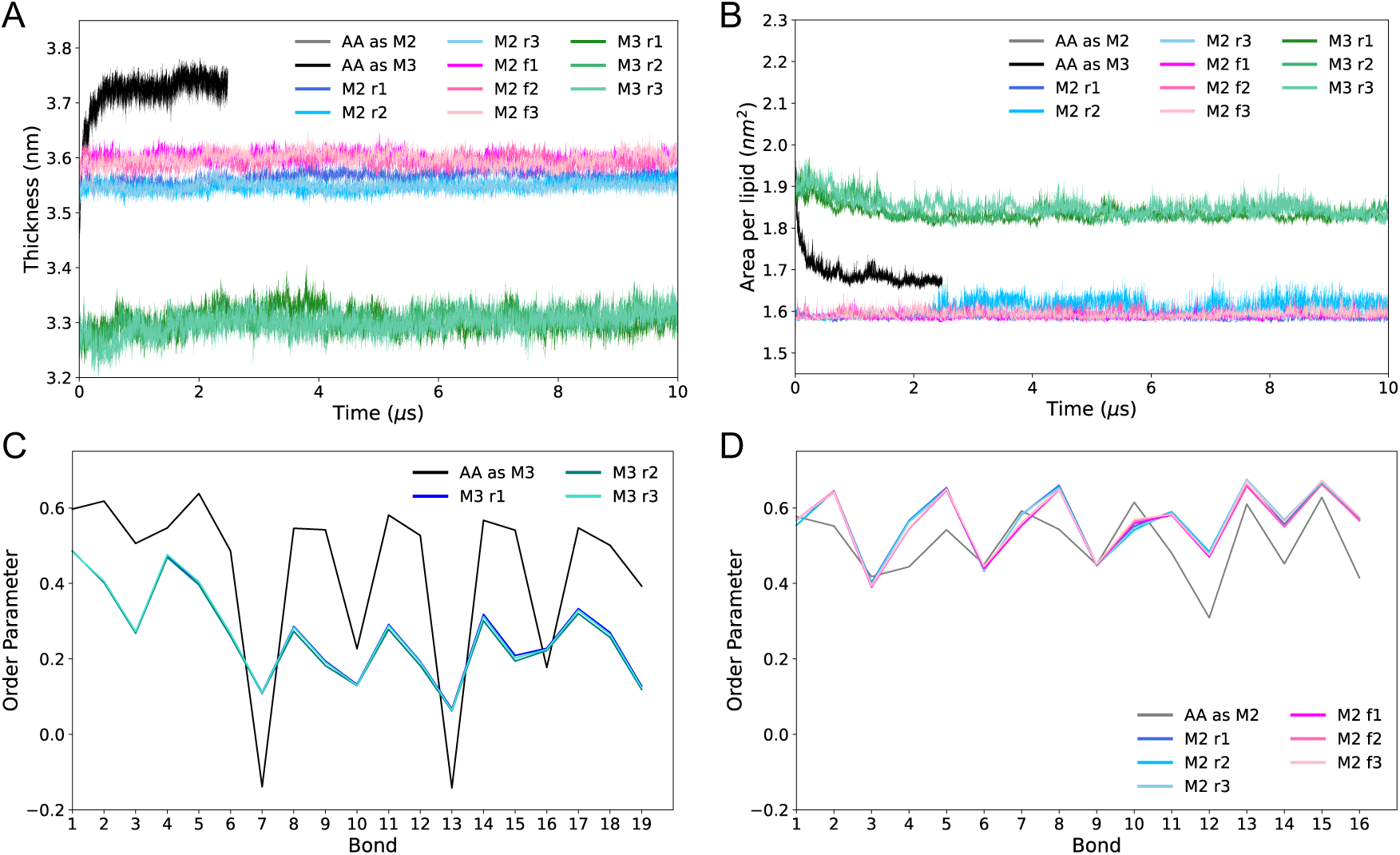
Physical properties of ReLPS in standard and fast Martini 2, and in Martini 3. **(A)** Membrane thickness, (the curves for AA as M2 and AA as M3 are almost completely indistinguishable) **(B)** area per lipid (the curves for AA as M2 and AA as M3 are almost completely indistinguishable), and **(C, D)** lipid order parameters of our new Martini 3 ReLPS model and for the Martini 2 standard and fast ReLPS model respectively. The plots show data taken from 3 simulation replicas with the Martini 2 standard (M2 r1-3), Martini 2 fast (M2 f1-3), Martini 3 (M3 r1-3) models, and atomistic simulations mapped to Martini 2 (AA as M2) or mapped as Martini 3 (AA as M3).

### 3.3 Testing the permeability of ReLPS with octane, DPC, DOPC and PMB1

Having achieved increased mobility in the fast ReLPS model we next sought to characterise the permeability of membranes containing these modified lipids and to compare them with the standard Martini 2 and the newly parameterised Martini 3 models. The concern being that the greater mobility may render the membranes too permeable. To this end we explored the permeation of DPC, octane and the antibiotic polymyxin B1 (PMB1) *via* equilibrium MD simulations. PMB1 is used as a permeabilising agent for LPS-containing membranes but has previously been shown (*via* atomistic and CGMD) simulations to be susceptible to kinetic barriers to permeation ^6–8^.

We used membrane models with 100% of either Martini 2, Martini 2 fast or Martini 3 ReLPS in the outer leaflet and 100 % PL in the inner leaflet. Octane was able to insert easily into the Martini 2 models, with all 40 octane molecules (4 molecules in 10 simulation replicas) inserting into both the standard and fast Martini 2 membrane models. In contrast, in the Martini 3 containing membrane, only 4 out of 40 octane molecules inserted. A similar trend is observed for DPC, but in this case we did not observe any penetration into the Martini 3-containing membrane at all. This is shown quantitatively by measuring the number density of each system components as a function of position along the Z dimension (perpendicular to the membrane plane) (**fig 4**). We also tested the ability of a phospholipid; DOPC, to insert into the LPS leaflet (10 x 10 µs each). Insertion of DOPC was observed in 8 out of 10 simulations of the standard Martini 2 model, in 7 out of 10 simulations of the fast model, and none in the Martini 3 model (**fig S17A).** This indicates that our fast ReLPS model, while increasing the lateral movement of lipids, does not render the bilayer over-susceptible to permeation by small molecules given the permeation behaviour is very similar to the Martini 2 model. Our data reveals the Martini 2 ReLPS gives the more expected insertion behaviour (compared to Martini 3)-we would expect a simple detergent molecule and a phospholipid to insert into the outer membrane. We note here that there are reports in the literature about oddities in membrane/lipid behaviour of Martini 3 for example that Martini 3 protein-water interactions need to be scaled down to successfully allow (i) phospholipid bilayer self-assembly around transmembrane protein dimers ^22^, and (ii) monomeric peptides to maintain a transmembrane state ^28^. In other words, the protein-water interactions had to be made less polar to enable insertion of the protein (Glycophorin A). This is consistent with our findings here that DOPC and DPC do not penetrate into the Martini 3 ReLPS-PL bilayers, because they prefer to be in the aqueous region, i.e. the water-hydrophobic tail interaction is too attractive, just as reported for Glycophorin A by Wade and co-workers ^22^. This also explains why water penetrates further into the membrane core in the Martini 3 simulations compared to Martini 2 - the water-hydrophobic tail interaction is again too attractive. We show this semi-quantitatively using steered MD (**fig S18**). Greater force is required to pull DPC into the core of a Martini 3 ReLPS-PL bilayer compared to the Martini 2 versions, where a barrier is experienced by DPC when entering the hydrophobic core of the membrane.

**Figure 4.**
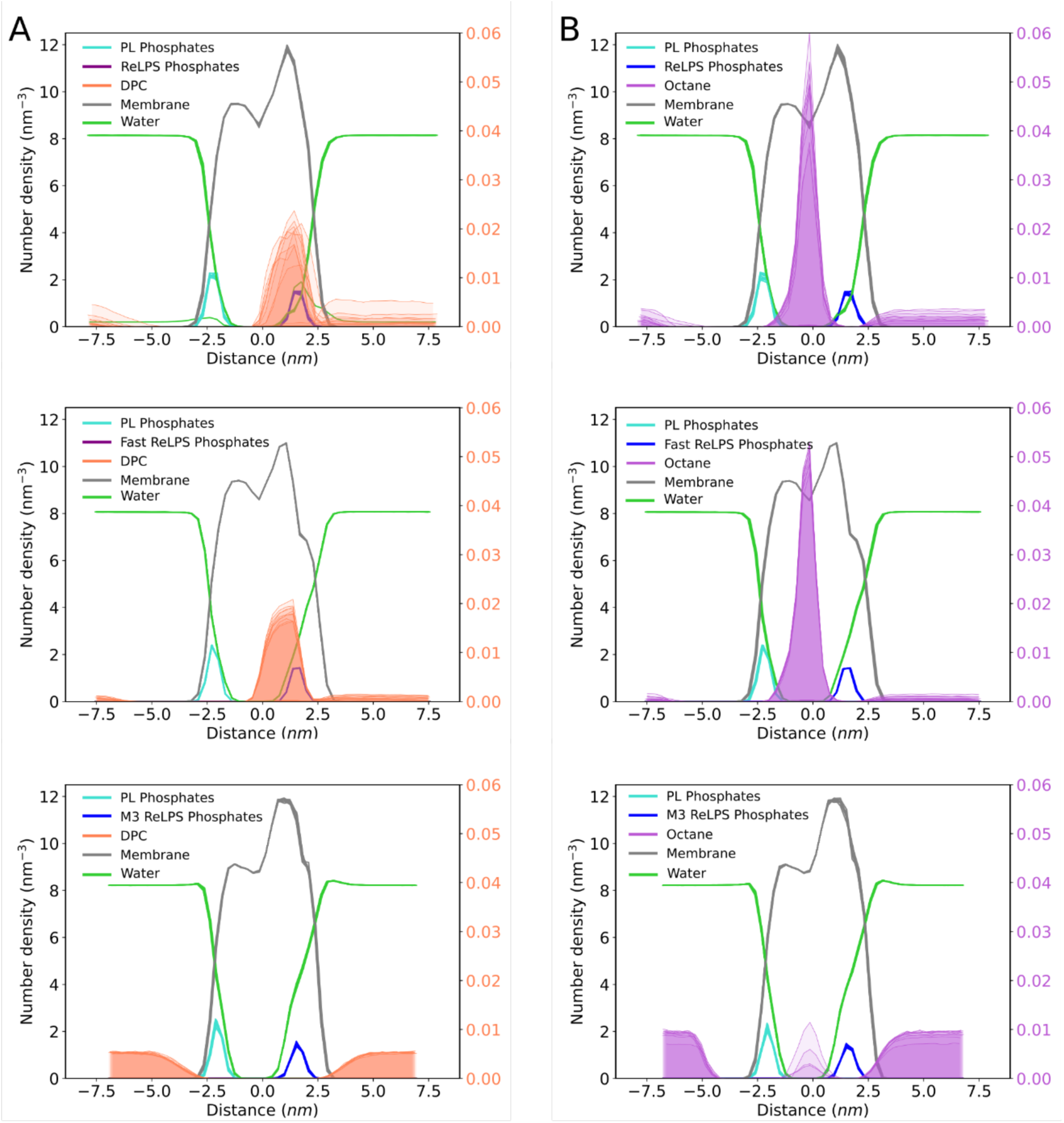
Permeability of ReLPS to DPC and octane. Number densities computed along the bilayer normal are shown in transparent surface for DPC **(A)** and octane **(B)** in standard Martini 2 (top), fast Martini 2 (middle), and our Martini 3 model (bottom). In each of the systems containing DPC, one DPC molecule was simulated. In each of the systems containing octane, 4 octane molecules were simulated. Number densities for the phosphates of the inner leaflet (PL Phosphates) or outer leaflet (ReLPS Phosphates), and all the lipid molecules in the membrane (Membrane) are also depicted to aid the analysis. Each plot shows data computed for 10 independent 10 μs-long simulation replicas. The axes labels on the right correspond to the number densities from DPC (A) or octane (B), a different scale was used given the much smaller numbers compared to the other system components.

We next tested the penetration of polymyxin B1 (PMB1), a last resort lipoprotein antibiotic, into the ReLPS leaflet in membranes modelled with standard and fast Martini 2. While we did not observe full penetration into the ReLPS-containing leaflet of the membrane with either model, we observed deeper penetration of PMB1 with fast ReLPS compared to standard Martini 2 (**fig 5A, 5B, S17B, S17C**). In addition, partial density profiles along the axis perpendicular to the plane of the membrane shows a greater density of PMB1 molecules embedded within the ReLPS monolayer in the fast ReLPS model. Specifically, in the fast ReLPS models, a greater number of PMB1 molecules are located at the lipid headgroup/tail interface whereas with standard ReLPS the PMB1 are located largely at the water-headgroup interface; adhering to the membrane but not embedded. (**fig 5C, D**). The fact that PMB1 molecules do not fully insert into the fast ReLPS leaflet provides further evidence that this model does not cause the bilayer to be over-permeable. The absence of a Martini 3 model of PMB1 precluded the insertion study of that model.

**Figure 5.**
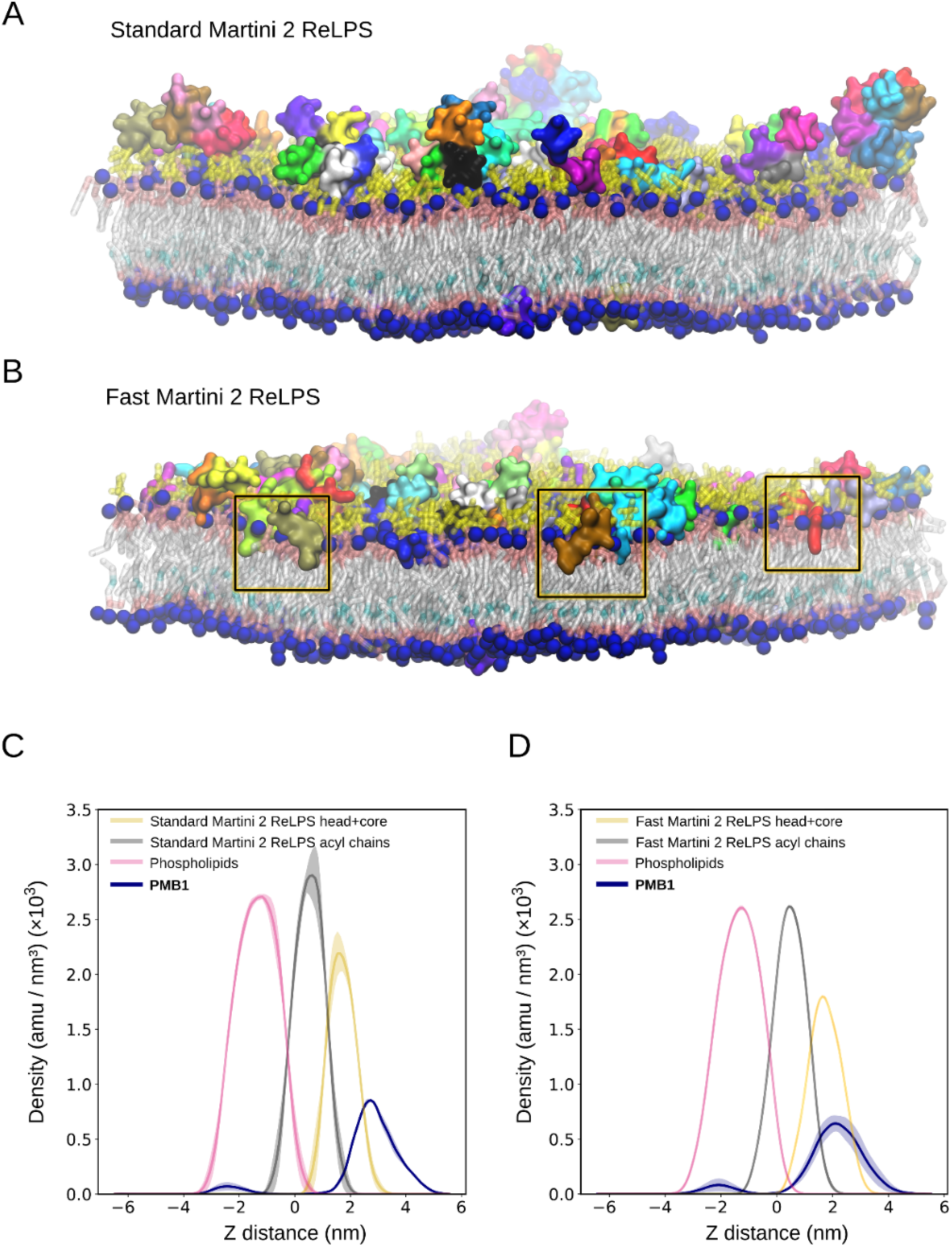
Interactions of PMB1 beads with standard and fast Martini 2 ReLPS. **(A)** Molecular representations of PMB1 association with the OM model containing standard Martini 2 ReLPS and **(B)** fast Martini 2 ReLPS in the outer leaflet, at 5 μs simulation time. In **(C)** standard Martini 2 and **(D)** fast Martini 2 ReLPS-containing outer membrane models, the densities of the PMB1 are shown relative to the respective ReLPS heads, tails, and of phospholipids along the Z position of the membrane.

### 3.4 Outer membrane protein-ReLPS interactions

Perhaps a more stringent test of the predictive properties of our ReLPS models is their ability to identify known ReLPS binding sites on *E. coli* outer membrane proteins (OMPs). Therefore, we simulated the trimeric porin OmpF using the X-ray structure determined at 1.98 Å (PDB:3K1B) using Martini 2 (standard and fast) and Martini 3. OmpF was embedded in a membrane containing only 2% ReLPS and 98% phospholipids in the outer leaflet to ensure accelerated lateral motion of ReLPS over 3 x 4 µs simulations.

We calculated the residence time i.e. the lifetime of ReLPS contacts with individual OmpF residues using PyLipID ^29^. This showed good agreement with the experimentally known LPS binding sites from two independent studies (**fig 6A**), one in which LPS-binding OmpF residues were proposed based on mutagenesis experiments and on the X-ray structure of the homologous protein OmpE ^30^, and the other from cross-linking studies ^26^. The mutagenesis experiments showed that mutations of positively charged residues (K25Q, K160Q, K209Q, K210Q, R235Q, K253Q, K277Q, K279Q, K281Q, K323Q) reduced the amount of LPS bound to OmpF, whereas crosslinking experiments highlighted V155, V174, G176, T216 and L259 (referred to as V177, V196, G198, T238, L281 by Webby *et al.*) to be involved in LPS binding. Our simulations, across all Martini models of ReLPS, were able to reproduce these data and further quantify these interactions with OmpF thereby suggesting ‘hotspots’ for LPS binding.

**Figure 6.**
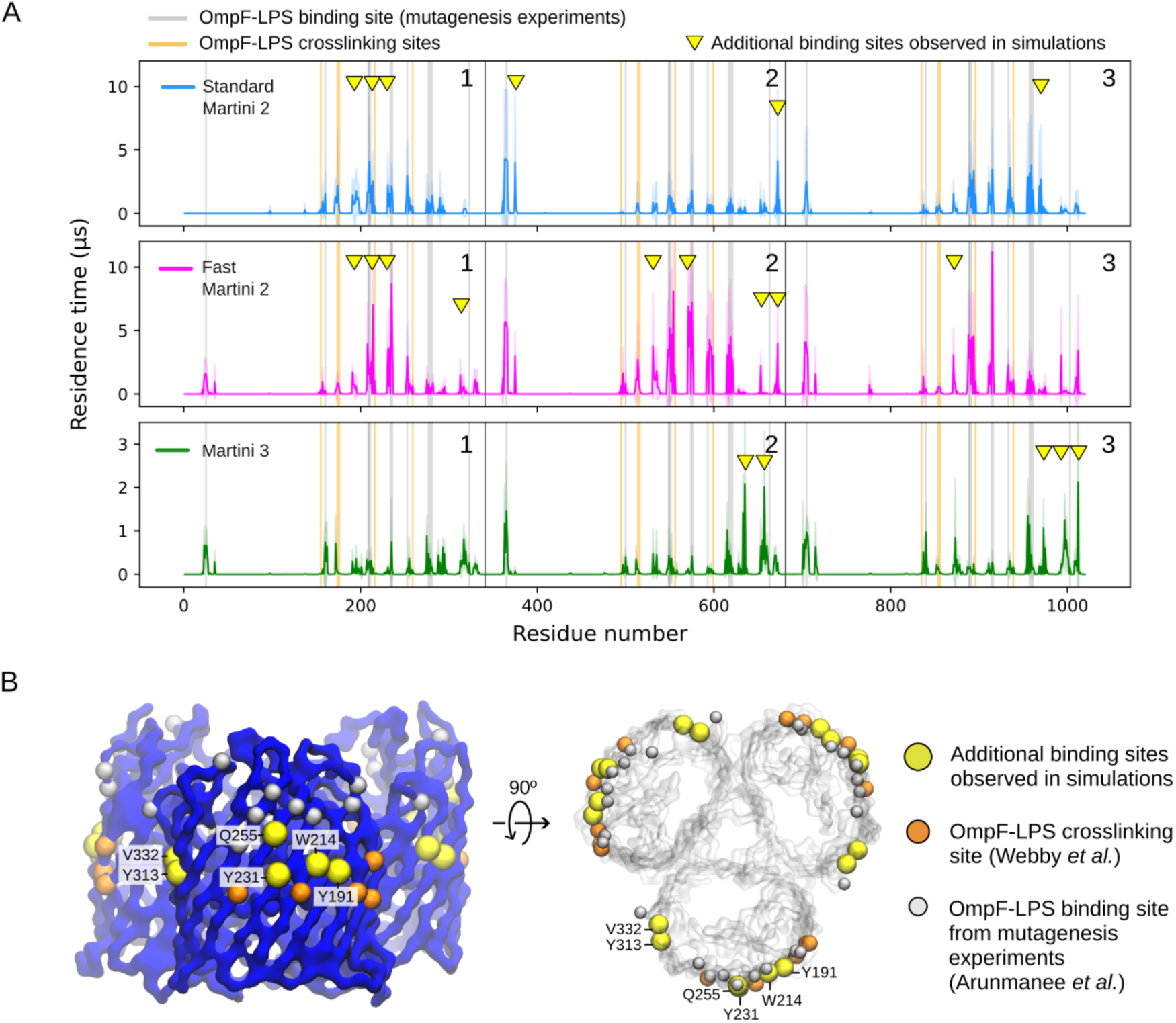
Residence times of fast and standard Martini 2 ReLPS, and our Martini 3 ReLPS with OmpF compared with experimental data on LPS binding sites of OmpF. **(A)** Standard (blue) and fast Martini 2 ReLPS (magenta), and our Martini 3 ReLPS model (green). Grey lines indicate the residues that are proposed to contribute to LPS binding by mutagenesis experiments ^30^. Orange lines indicate OmpF-LPS crosslinking sites ^26^. The yellow triangles indicate newly identified OmpF residues from simulations whose residence times with ReLPS are comparable to or larger than those that align with experimental data. The y-axis for the Martini 3 model is magnified for visual clarity of its residence time peaks. **(B)** A side view (left) and a top view of OmpF (right) with the newly identified residues shown as yellow spheres and labelled.

In addition to these simulations revealing ReLPS binding sites in agreement with experimentally determined LPS binding sites, we consistently observed other residence time peaks non-uniformly on all the OmpF monomeric subunits with all the CG models (**fig 6A**). We note that the pattern of interactions differs between OmpF monomers due to the low concentration of ReLPS used in the simulations. On the other hand, a higher number of ReLPS molecules in the leaflet could lead to LPS-LPS aggregation and reduce sampling protein-LPS interactions. We observed that the fast Martini 2 ReLPS model showed highest residence time peaks on average followed by the standard Martini 2 model and then the Martini 3 model (note the magnified y-axis for Martini 3 in **fig 6A**). Our fast model exhibits higher residence time compared to the standard model as it experiences less LPS-LPS aggregation in the membrane before it associates with the protein due to the scaled-down LPS-LPS interactions. Therefore, compared to the standard model, it forms contacts with OmpF at an earlier point in time. Further, upon association of a fast ReLPS molecule with OmpF, the contacts between its 6 acyl chains and OmpF (whose non-bonded potentials have not been scaled at all) form long-lasting interactions similar to the standard model.

To aid in visualising the residence times of the three Martini models, we fitted the coordinates of the OmpF backbone along the x-y plane and plotted the density of the center of mass of ReLPS molecules for each individual simulation (6 x 12 µs) (**fig S19B**). The less localised density of the Martini 3 ReLPS molecules around the OmpF trimer clarified that they interacted with the protein but did not stay attached for as long as the Martini 2 models. Nonetheless, similar to the fast Martini 2 model, our Martini 3 model did explore and interact with a larger surface area of the protein compared to the standard Martini 2 model which turned out to be restricted to certain areas on the protein once bound.

To assess the overall spatial localisation of ReLPS with respect to the OmpF trimer, we computed their radial distribution functions (RDFs) with reference to the OmpF trimer and observed an enrichment of fast Martini 2 ReLPS compared to the standard model (i.e. the first peak, although located in the same position in the fast model, is twice than that of the standard model, **fig S19A**).

We then sought to do a simple contact analysis between OmpF residues and ReLPS molecules, which matched the residence time peaks and confirmed that our simulations not only reproduced experimentally determined LPS binding sites but also predicted additional sites on OmpF that form favourable interactions with ReLPS, specifically residues Y191, W214, Y231, Q255, Y313, V332. (**fig S20**). The locations of these residues are shown on the OmpF backbone structure alongside those of LPS binding sites suggested by experiments in **fig 6B**.

In addition, we calculated the number of contacts versus time of all the ReLPS models with OmpF (**fig S21, S22**) across all simulations to check if protein-ReLPS interactions are dynamic, which is essential to ensure the interactions we observe are not dependent on the initial system configuration. From this data, we calculated the number of dissociation events from OmpF of each ReLPS molecule in each simulation replica across all Martini models. To do this, we considered an ReLPS to be in contact with OmpF when there were ≥ 30 pairs of contacts between OmpF beads and ReLPS beads, dissociation was then deemed to occur when there were zero contacts between the two. Taking into consideration all simulation replicas and all ReLPS molecules in each of them, we determined that our Martini 3, fast Martini 2, and the standard Martini 2 models undergo a total of 1029, 43, and 67 dissociations from OmpF respectively. This is consistent with residence time **(fig 6),** RDF and 2D density (**fig S19)**, number of contacts per OmpF residue **(fig S20)**, and number of protein-ReLPS contacts versus time **(fig S21 - S22**) analyses across all Martini models of ReLPS, which indicate that our Martini 3 model interacts with the protein more transiently compared to the Martini 2 models at this LPS concentration. Nonetheless, all Martini ReLPS models are able to dynamically bind OmpF throughout the trajectory and reproduce experimental LPS-binding sites on the surface of OmpF.

We next performed simulations of outer membrane systems containing 7 x OmpF trimers (100% ReLPS in the outer leaflet) with standard and fast Martini 2 ReLPS (**fig 7A, B** respectively). In these simulations, fast ReLPS molecules that were initially in contact with OmpF explored larger membrane surface area compared to its standard Martini counterpart. Not only did fast ReLPS diffuse more, but we also observed multiple instances of ReLPS-OmpF dissociation. Moreover, global ReLPS mixing was increased in the fast ReLPS simulations compared to those containing standard ReLPS, as revealed by tracking the lateral movement of one phosphate bead (PO1) of each individual ReLPS molecule.

**Figure 7.**
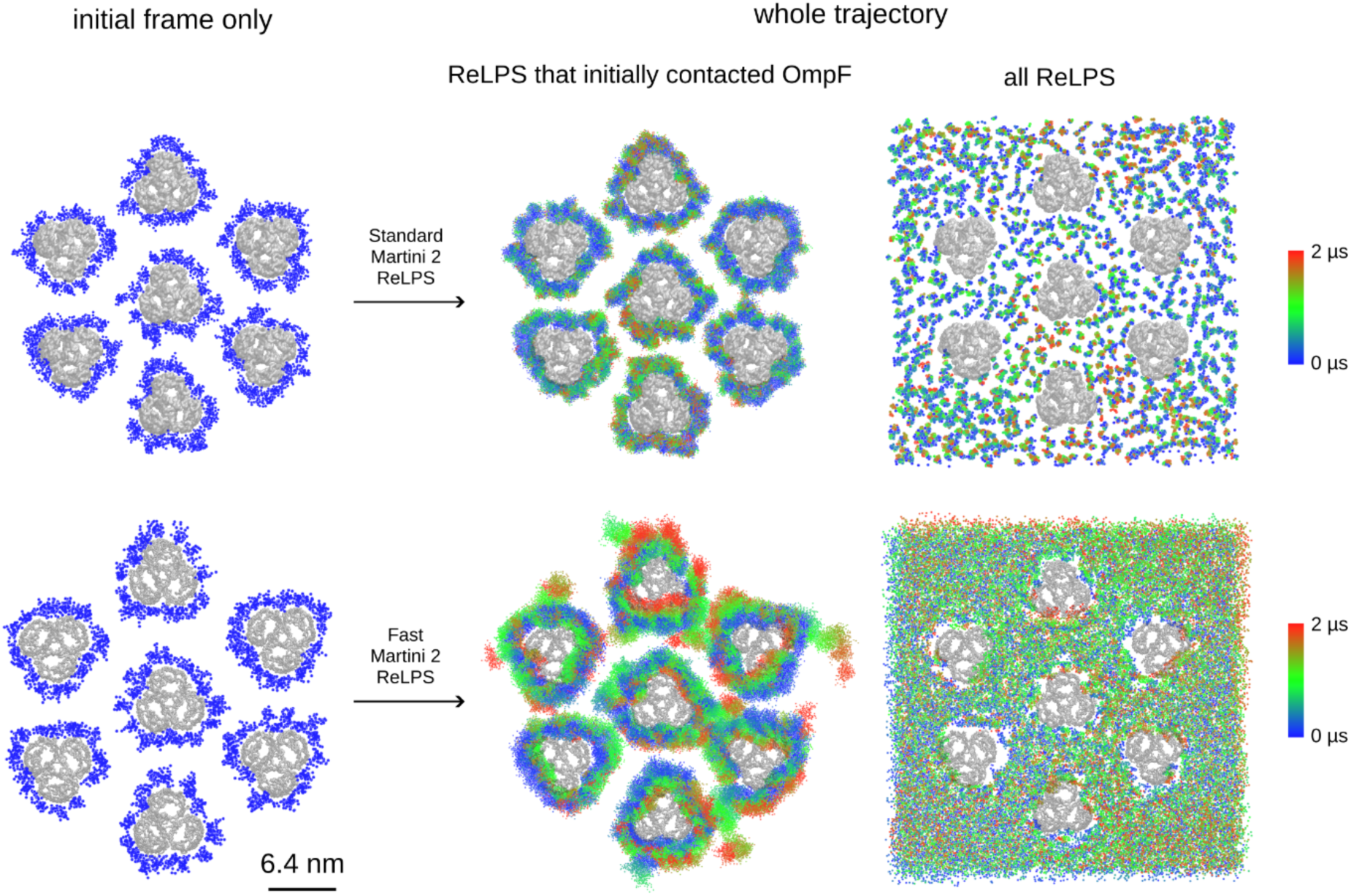
ReLPS lateral mobility in an outer membrane model system containing 7 OmpF trimers. Individual ReLPS molecules were tracked along the full trajectory (2 μs) after protein fitting. (A) shows the mobility of standard Martini 2 ReLPS, whereas (B) is the same as (A) but for fast Martini 2. The middle image in each row shows the respective movement of ReLPS that were initially in the vicinity of the protein. The right panel shows the tracked path of one bead of each ReLPS in the systems, coloured by time clearly showing that fast ReLPS has increased mixing rate.

### 3.5 Distribution of OMPs within the outer membrane

We have recently reported simulations of a patch of an ‘OMP island’ as observed in AFM images using standard Martini 2 RaLPS. Our simulated model, whose near neighbour distance was consistent with AFM data ^31^, provided the most realistic molecular simulation model of an *E. coli* OMP island to date ^26^. This model contained 7 different OMPs including multimeric proteins such as the BAM complex (in the ratio OmpF:BtuB:FhuA:FhuE:FepA:LptDE:BAM=102:20:20:19:19:19:19) giving an OMP island model spanning 150 nm^2^ along the membrane plane (**fig 8A**). Here this OMP island provides a useful test of the ability of our new ReLPS models to maintain the spatial arrangement of OMPs.

**Figure 8.**
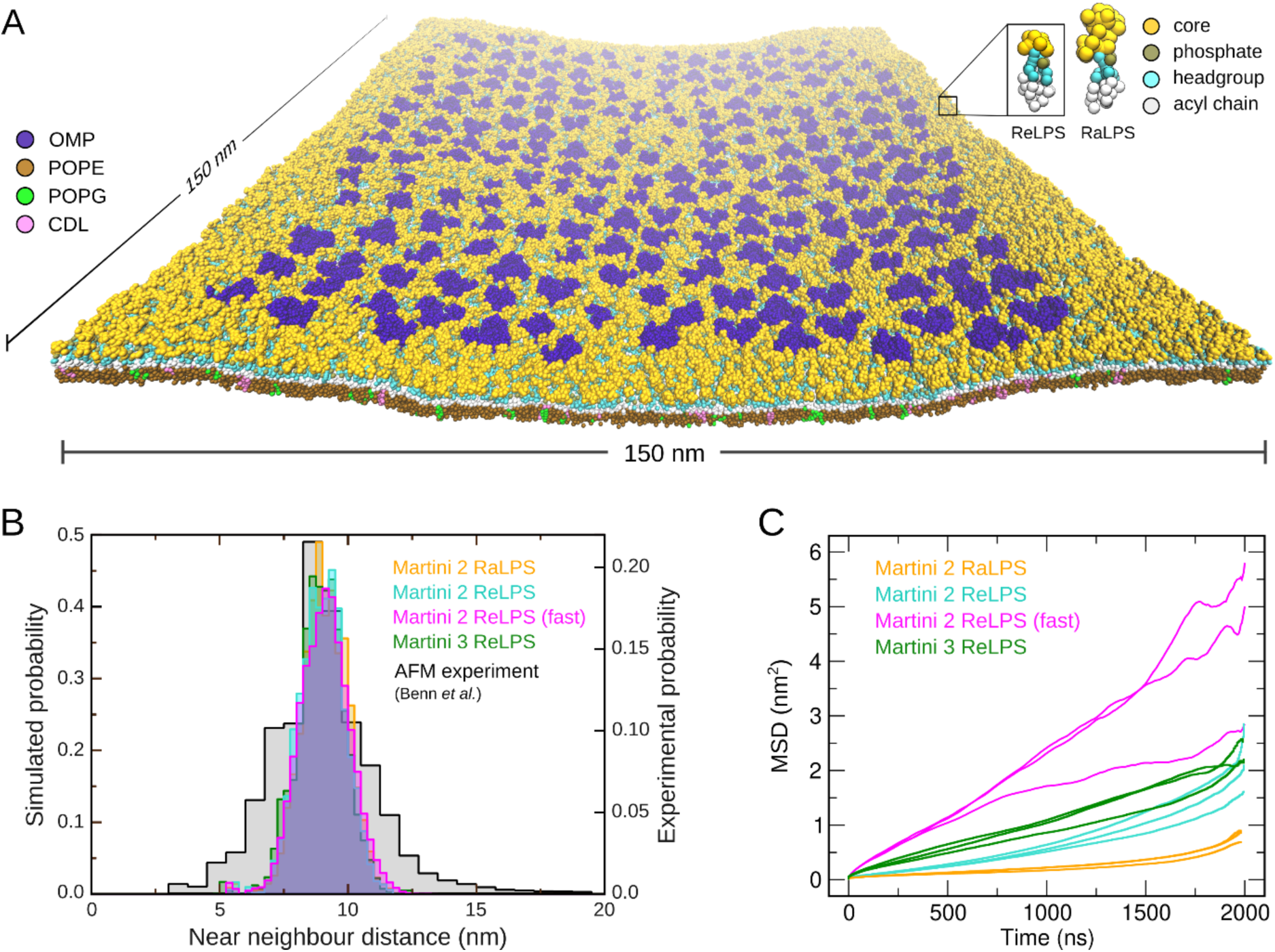
Distribution of OMPs within outer membrane models containing standard or fast ReLPS models from Martini 2 and 3, and the standard Martini 2 RaLPS model. **(A)** A snapshot of the OMP island embedded in a Martini 2 ReLPS-containing outer membrane model. Near-neighbour distances were calculated from this OMP island model. **(B)** Near-neighbour distance distribution of OMPs shown as a histogram to enable comparison between our simulated ReLPS models and with AFM data. The AFM data is based on experiments conducted by Benn et al ^31^. **(C)** Lateral MSD of OMPs are plotted for all three individual replicas of each simulated system. For reference, the MSD of the ReLPS molecules in the OMP island simulations from each of the three ReLPS models is provided in **fig S23.**

We simulated four new systems, in which the RaLPS from our original work ^26^, was replaced with the standard Martini 2 ReLPS, the fast Martini 2 ReLPS, and Martini 3 ReLPS models. Note that RaLPS has a longer glycan core (39 beads) compared to that of ReLPS (10 beads) (shown in **fig 8A**). The distribution of near-neighbour distances of OMPs in the simulated OMP island (3 x 2 μs simulations of each system) was maintained, showing good agreement with the AFM data and the original previously reported Martini 2 RaLPS simulations (**fig 8B**). Furthermore, the near-neighbour distance distribution between the different ReLPS models also agree, confirming that our fast Martini 2 and new Martini 3 ReLPS models are able to reproduce the behaviour of the standard Martini 2 model even in much larger systems. Meanwhile, by calculating the MSDs of OMPs, we show that the OMPs achieved varied magnitudes of lateral diffusion between the different ReLPS models. Here, we observed that the MSD of proteins embedded within the membrane containing fast ReLPS is enhanced (**fig 8C**) while our fast Martini 2 ReLPS model itself has a greater MSD compared to the other Martini LPS models included in this study (**fig S23**). The lowest MSD of OMPs is when they are in the Martini 2 RaLPS containing OM model, which is unsurprising given (1) the OMPs are able to form a greater number of interactions with RaLPS molecules compared to the ReLPS models, owing to the longer length of the oligosaccharides in the former and (2) the MSD of RaLPS itself is lower than the ReLPS model (**fig S23**).

We note here that the distribution of near neighbour distances is wider in the AFM data **(figure 8 (B))** compared to the distributions from simulations likely due to the AFM images being taken from the surface of an entire cell, whereas the simulations only incorporate a single ‘OMP island’ albeit containing 237 OMPs. Overall, our data show that our new ReLPS models are able to maintain experimentally determined OMP distributions within LPS-containing outer membrane models.

## 4 DISCUSSION

Here we have sought to address the need for faster kinetics within LPS-containing model membranes as well as to complete the Martini 3 *E. coli* LPS modelling toolkit. Our newly parameterised Martini 3 ReLPS model is compatible with existing Martini 3 models of other lengths of LPS molecules, and we note here that there is no reason why it cannot be used in combination with other models for simulations of LPS molecules of varying lengths, despite our model using a slightly different mapping scheme ^16^. Overall, our Martini 3 ReLPS shows agreement with its atomistic and Martini 2 counterparts. We note that there is some deviation in the area per lipid and membrane thickness of the Martini 3 ReLPS model from the Martini 2 versions, while some of these differences may arise due to general differences between the two force-fields, further work is needed to fully ascertain the origins of these differences and whether they have any functional consequences. There is greater water penetration towards the bilayer core in in the Martini 3 ReLPS bilayer (and this better matches all atom behaviour), which may cause the larger APL values for the Martini 3 model, as the water interdigitates more between the lipid headgroups. It is also of note that these differences are within 5% and 9% of the atomistic APL reference value for the Martini 2 models and Martini 3 model respectively (obtained from the coarse-grained pseudo-trajectory, i.e. the atomistic trajectory mapped to their respective Martini model). These deviation ranges are similar to previously reported values for the original Martini 2 Model (4-5%) ^32^ and similar Lipid A or LPS molecules reported by Vaiwala *et al*. ^16^ (2-4%), albeit the calculation of the APL in the latter was computed in a different manner i.e. dividing the surface of the membrane by the amount of lipids present in one leaflet.

Our fast Martini 2 model is stable over long simulations not only in symmetric bilayers but also in more complex model membranes, including in the presence of embedded proteins. Furthermore, this model not only reproduces the biophysical properties of the standard model very well (membrane thickness, APL, lipid tail order parameters as shown in **fig 3**), but it also shows enhanced lateral diffusion as well as greater conformational freedom within individual molecules. The fast model compared to its standard Martini 2 counterpart is slightly more permeable to polymyxin B1, an antimicrobial peptide known to be able to localise into the outer membrane of *E. coli*. We have also demonstrated that the fast model is not overly permeable to water and ions as clearly observed by the lateral partial number density data (**fig S12**). One key advantage of this fast Martini 2 ReLPS model is that by increasing ReLPS mobility, (i) the relative orientation of the proteins embedded in the membrane and (ii) the patterns of protein-lipid interactions are not strongly dependent on the initial configuration.

It is useful to reflect on potential limitations. Overall, we note here that the models for fast Martini 2 ReLPS and Martini 3 ReLPS could both benefit from further testing by the computational community. The issues of penetration of hydrophobic entities into Martini 3 membranes (reported here for ReLPS-PL bilayers and also by others for phospholipid-only containing bilayers ^22^) preclude the development of a fast Martini 3 ReLPS model, until these are resolved. We posit that the straight-forward usage of these models within the GROMACS package is a clear benefit for potential users. However, the user should carefully consider the scientific question they are addressing when using the fast ReLPS model, since its molecular chemistry is slightly altered due to the scaling of its original charges. While we are aware that algorithmic enhanced sampling methods such as temperature replica exchange molecular dynamics (T-REMD) can be used to extend the sampling of phase space even in the NPT ensemble, for LPS they would require impractical computational resources, and as we have previously shown *via* Hamiltonian replica exchange, LPS mixing requires long simulation times to potentially converge ^5^. Therefore, despite its tweaked chemistry our model offers a practical and ‘off the shelf’ solution for simulations of ReLPS-containing bilayers.

## Supporting information

additional text within the supplementary information

## TOC graphic

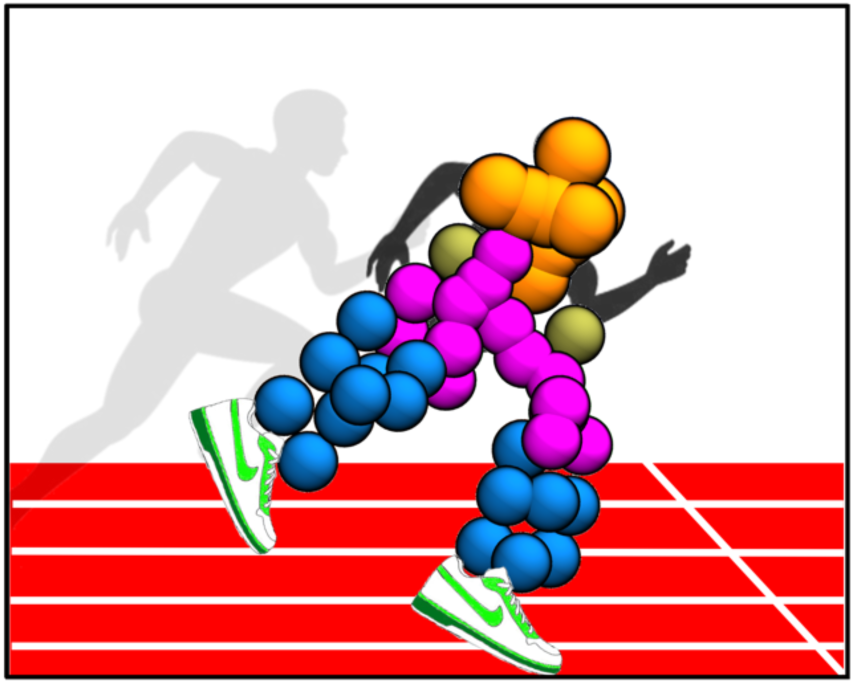

