## Supplementary material for "Faster but not sweeter: A model of *E. coli* Re-level lipopolysaccharide for Martini 3 and a Martini 2 version with accelerated kinetics": S1-S3

1. Department of Biochemistry,
   University of Oxford
   Oxford
   OX1 3QU

2. Departamento de Física Aplicada,
   Universidade de Santiago de Compostela,
   Spain

3. Departamento de Química Orgánica,
   Universidade de Santiago de Compostela,
   Spain

4. MD.USE Innovations S.L.,
   Edificio Emprendia, 15782 Santiago de Compostela,
   Spain

* These authors contributed equally

### 1. Supplementary Methods

#### 1.1. Details for setting up the atomistic (CHARMM36m) simulation

NPT conditions were maintained using a Nosé-Hoover thermostat [1,2], and a Parrinello-Rahman barostat [3] with a compressibility of $4.5 \times 10^{-5}$ $bar^{-1}$. The integration timestep was set to 2 fs, and bonds to hydrogens were constrained with the LINCS algorithm [4]. Long-range electrostatics were computed using particle-mesh Ewald [5] with a tolerance of $10^{-6}$, 4th-order spline interpolation, and a maximal mesh size of 0.12 nm; van der Waals interactions were shifted smoothly to zero between 1.0 and 1.2 nm.

#### 1.2. Details for setting up coarse-grained (Martini 2 and 3) simulations

All the systems were minimised using the steepest descent algorithm until the forces converged to less than 1000 kJ/mol/nm. With the Martini 2 force field, the systems were equilibrated once at 313 K using a 20 fs timestep, except for the PMB1 simulations where a 10 fs timestep was used. The time of equilibration for simulations of the OMP island and PMB1 were 30 ns and 100 ns respectively. All other Martini 2 systems were equilibrated for 5 ns each. During equilibration, the cut-off algorithm combined with the potential-shift modifier was employed to treat both van der Waals and coulombic interactions. The van der Waals and coulombic radius was set to 1.1 nm. The pressure was maintained at 1 bar with semi-isotropic pressure coupling using the Berendsen barostat [6], the compressibility was set to 3.0-4 /bar, the temperature was maintained at 313 K using the V-rescale thermostat [7] and a time constant of 1 ps.

With the Martini 3 force field, systems were equilibrated step-wise using the Berendsen barostat for a total time of 150.6 ns i.e. (1) with 1 fs timestep at 200 K for 50 ps, (2) with 1 fs timestep at 313 K for 50 ps, (3) with 5 fs timestep at 313 K for 500 ps, (4) with 10 fs timestep at 313 K for 50 ns, (5) with 10 fs timestep at 313 K for 100 ns. From step 3 to step 5, the protein backbone and ReLPS phosphate beads were position restrained with 50 kJ/mol/nm harmonic potential. For equilibration and production stages, the reaction-field algorithm was used to treat coulombic interactions and the cutoff with the potential-shift modifier algorithms to treat van der Waals interactions in Martini 3 simulations. For the membrane permeability simulations, the octane/ DPC/ DOPC molecule was position restrained during equilibration. For steered molecular dynamics (MD) simulations, this equilibrated snapshot was used as the initial frame and a force of 1000 kJ/mol/nm was applied on the center of mass of DPC in the -Z direction to pull it from water into the bilayer using at the rate of 1 nm/ns. These steered MD simulations were performed in 5 replicas using the standard Martini 2 and Martini 3 ReLPS models. All production simulations including the steered MD simulations used the Parrinello-Rahman barostat [3] along with the velocity rescaling thermostat [8].

#### 1.3. Simulation analysis and visualisation

The GROMACS package [9] version 2019.4 or higher was used to setup, run, and post-process all simulations. PyLipID [10] was used to calculate residence times between ReLPS and outer membrane proteins (OMPs) using a lower and upper cutoff of 0.485 and 0.8 nm to define contacts. RMSD, distances, lateral diffusion, density maps, radial distribution functions and contacts were analysed using GROMACS analysis tools: gmx rms, gmx distance, gmx msd (-lateral z), gmx densmap, gmx rdf, gmx mindist respectively. The lateral diffusion coefficients of the Martini 2 ReLPS models were obtained for independent 1 μs-long segments from a linear fit of the MSD using a time window of 20-100 ns for the fit. Area per lipid (APL) and membrane thickness were analysed using FATSLiM [11]. Phosphate moieties were used to define the group for leaflet identification. Lipid order parameters were calculated using a modified version of the do-order-multi.py script available on the www.cgmartini.nl website (http://www.cgmartini.nl/index.php/downloads/tools/229-do-order). VMD was used for visualisation of all systems [12]. NumPy [13] was used during postprocessing and analyses of data. Matplotlib [14] and Xmgrace (https://plasma- humphreygate.weizmann.ac.il/Grace/) were used for plotting. The near-neighbour (NN) index [15] was calculated using the following equation:

$$NN\ Index = (unique\ neighbour\ number - N_{NN}) / (N_{LPS} - 1 - N_{NN})$$

where unique neighbour number refers to the number of unique LPS molecules contacted by the reference LPS throughout the trajectory, $N_{NN}$ is the number of surrounding LPS molecules in a single frame of the trajectory (taken as 6), $N_{LPS}$ is the number of LPS molecules in the simulation.

##### 1.4. Details of obtaining the fast ReLPS Martini 2 model

We first investigated reduction of the magnitude of the charges (by 100, 50 and 25 %) on all anionic beads of ReLPS. Each of these settings were tested with neutralising monovalent ($Na^+$) and divalent ($Ca^{2+}$) cations. To quantify the impact of these alterations to charges we consider the following metrics: mean squared displacement (MSD), nearest neighbour index (NNI), APL, and partial densities. The NNI provides a measure of the number of neighbours encountered by an ReLPS molecule throughout a trajectory and thus gives an indication of lipid mixing. Overall, from the MSD (**fig S6A**) and NNI data (**fig S6B**), compared to the standard Martini 2 system neutralised by $Ca^{2+}$ ions (labelled as 'Q100%, $Ca^{2+}$'), we observed that individual ReLPS molecules achieved greatest lateral motion when the charge of ReLPS anionic beads and cations were scaled down by 50% (labelled "Q50%, $Na^+$").

We then investigated why there is a maximum in the curves for MSD and NNI of ReLPS as a function of charge (**fig S6A, B**). In **fig S7A**, if we consider the RDF curves of ReLPS phosphates (PO)-$Ca^{2+}$, as the PO bead charge is reduced, there are fewer and fewer PO and $Ca^{2+}$ in proximity with each other. Concomitantly, the PO-PO RDF curves reveal the reverse trend with decreasing charge. As more and more POs come closer to each other, they repel due to electrostatics leading to increased MSD and NNI. However, at the point at which the charge is totally removed from the PO bead, despite their proximity, there is no electrostatic repulsion between PO-PO beads and therefore the average MSD and NNI drop slightly. We provide the non-bonded potential plots for each charge regime to further highlight this point (**fig S7A**). The same trend is observed with PO-$Na^+$ interactions (**fig S7B**).

The partial density profiles of various components of this charge-modified (Q50%, $Na^+$) system suggest that the system is stable and comparable with those of the standard Martini 2 system (**fig S8A**). Concomitantly, there was a minimum increase in the APL from 1.67 ± 0.23 $nm^2$ to 1.73 ± 0.3 $nm^2$ (**fig S8B**). Therefore, based on these data, the model whereby the charges of both the ReLPS anionic beads and neutralising cations were scaled down by 50% was deemed to provide the best compromise between deviation from the standard Martini 2 model and faster kinetics, and was taken forward to the next step.

We next investigated scaling down the inter-ReLPS LJ interactions (which leads to a shallower well-depth of potential energy). We tested a variety of scaling combinations i.e. scaling the LJ interactions of only GlcN headgroup beads, only the Kdo core (S beads), and both GlcN and Kdo core beads simultaneously (**fig S9A-C**). The scaling regimes were evaluated by calculating MSD and NNI of ReLPS. These data revealed that scaling only the Kdo core had little impact on MSD and NNI, whereas greater impact was achieved when modifying either only the GlcN headgroup beads or GlcN and Kdo core beads. Thus, in the interests of limiting chemical modifications as much as possible, only alterations to the GlcN beads were considered hereon. Compared to the standard Martini 2 model, while scaling down the LJ interactions of the GlcN beads by 5 levels (as defined by Martini 2 conventions) led to the greatest lateral diffusion (**fig S9B**), we took into account the (1) change in the APL i.e. an increase less than 5% (**table S3**), (2) the partial density profiles showing a good agreement with the standard Martini 2 model (**fig S10A**), and (3) membrane thickness showing an increase (~1%) less than 0.1 nm (**fig S10B**). As a result, we considered the model with LJ scaled down by 2 levels.

#### 2. Supplementary Tables

**Table S1.** ReLPS CHARMM36m (all-atom) to Martini 3 (CG) mapping scheme.

| CG name | CG type | CHARMM36m names | | | | | | | | | | | | | | |
|---|---|---|---|---|---|---|---|---|---|---|---|---|---|---|---|---|
| ZM1 | SN6r | OA5 | HA5 | CA5 | CA6 | HA6S | HA6R | OA6 | | | | | | | | |
| ZM2 | TC1 | HA1 | CA1 | HA2 | CA2 | | | | | | | | | | | |
| ZM3 | SN6 | CA3 | CA4 | HA3 | HA4 | OA4 | HOA4 | | | | | | | | | |
| VS1 | TC4 | ZM1 | ZM2 | ZM3 | | | | | | | | | | | | |
| PO1 | D | OA1 | PA | OPA2 | OPA3 | OPA4 | | | | | | | | | | |
| ZM4 | N4a | OA3 | C31 | O31 | C32 | H32R | H32S | | | | | | | | | |
| ZM5 | P3 | NA2 | HNA2 | C11 | O11 | C12 | H12R | H12S | | | | | | | | |
| CF | P1 | C13 | H13R | O13 | HO13 | C14 | H14R | H14S | C15 | H15R | H15S | C16 | H16R | H16S | | |
| C1F | C1 | C17 | H17R | H17S | C18 | H18R | H18S | C19 | H19R | H19S | C110 | H110R | H110S | | | |
| C2F | C1 | C111 | H111R | H111S | C112 | H112R | H112S | C113 | H113R | H113S | C114 | H114R | H114S | H114T | | |
| CE | P1 | C33 | H33R | O33 | HO33 | C34 | H34R | H34S | C35 | H35R | H35S | C36 | H36R | H36S | | |
| C1E | C1 | C37 | H37R | H37S | C38 | H38R | H38S | C39 | H39R | H39S | C310 | H310R | H310S | | | |
| C2E | C1 | C311 | H311R | H311S | C312 | H312R | H312S | C313 | H313R | H313S | C314 | H314R | H314S | H314T | | |
| GM1 | TC1 | CB1 | CB2 | HB1 | HB2 | | | | | | | | | | | |
| GM2 | SN5r | OB5 | CB5 | CB6 | OB6 | HB6S | HB6R | HB5 | | | | | | | | |
| GM3 | TC1 | CB3 | CB4 | HB4 | HB3 | | | | | | | | | | | |
| VS2 | TC4 | GM1 | GM2 | GM3 | | | | | | | | | | | | |
| PO2 | D | OB4 | PB | OPB2 | OPB3 | OPB4 | | | | | | | | | | |
| GM4 | SN4a | OB3 | C71 | O71 | C72 | H72R | H72S | | | | | | | | | |
| GM5 | P3 | NB2 | HNB2 | C51 | O51 | C52 | H52R | H52S | | | | | | | | |
| CAB | N4a | C73 | H73R | O73 | C81 | O81 | | | | | | | | | | |
| C1A | C1 | C82 | H82R | H82S | C83 | H83R | H83S | C84 | H84R | H84S | C85 | H85R | H85S | | | |
| C2A | C1 | C86 | H86R | H86S | C87 | H87R | H87S | C88 | H88R | H88S | C89 | H89R | H89S | | | |
| C3A | C1 | C810 | H810R | H810S | C811 | H811R | H811S | C812 | H812R | H812S | C813 | H813R | H813S | C814 | H814R | H814S | H814T |
| C1B | SC1 | C74 | H74R | H74S | C75 | H75R | H75S | C76 | H76R | H76S | | | | | | |
| C2B | C1 | C77 | H77R | H77S | C78 | H78R | H78S | C79 | H79R | H79S | C710 | H710R | H710S | | | |
| C3B | C1 | C711 | H711R | H711S | C712 | H712R | H712S | C713 | H713R | H713S | C714 | H714R | H714S | H714T | | |
| CCD | N4a | C53 | H53R | O53 | C61 | O61 | | | | | | | | | | |
| C1C | SC1 | C62 | H62R | H62S | C63 | H63R | H63S | C64 | H64R | H64S | | | | | | |
| C2C | C1 | C65 | H65R | H65S | C66 | H66R | H66S | C67 | H67R | H67S | C68 | H68R | H68S | | | |
| C3C | C1 | C69 | H69R | H69S | C610 | H610R | H610S | C611 | H611R | H611S | C612 | H612R | H612S | H612T | | |
| C1D | SC1 | C54 | H54R | H54S | C55 | H55R | H55S | C56 | H56R | H56S | | | | | | |
| C2D | C1 | C57 | H57R | H57S | C58 | H58R | H58S | C59 | H59R | H59S | C510 | H510R | H510S | | | |
| C3D | C1 | C511 | H511R | H511S | C512 | H512R | H512S | C513 | H513R | H513S | C514 | H514R | H514S | H514T | | |
| KR1 | TC1 | CK2 | CK3 | HK31 | HK32 | | | | | | | | | | | |
| KR2 | TN2a | OK6 | CK6 | HK6 | | | | | | | | | | | | |
| KR3 | SP1r | CK4 | HK4 | OK4 | CK5 | HK5 | HKO5 | OK5 | | | | | | | | |
| VS3 | TC4 | KR1 | KR2 | KR3 | | | | | | | | | | | | |
| KS1 | SQ5n | CK1 | OK11 | OK12 | | | | | | | | | | | | |
| KS2 | SP3 | CK7 | HK7 | OK7 | HKO7 | CK8 | HK81 | HK82 | OK8 | HKO8 | | | | | | |
| KR4 | TC2 | CL2 | CL3 | HL31 | HL32 | | | | | | | | | | | |
| KR5 | TN2a | OL6 | CL6 | HL6 | | | | | | | | | | | | |
| KR6 | SP4r | CL4 | CL5 | HL4 | OL4 | HLO4 | HLO5 | HL5 | OL5 | | | | | | | |
| VS4 | TC4 | KR4 | KR5 | KR6 | | | | | | | | | | | | |
| KS3 | SQ5n | CL1 | OL11 | OL12 | | | | | | | | | | | | |
| KS4 | SP4r | CL7 | HL7 | CL8 | HL81 | HL82 | OL8 | HLO8 | HLO7 | OL7 | | | | | | |

**Table S2.** Martini 3 ReLPS bonded parameters

Table S2A Bond potential parameters

| I | j | $b_{eq}$ (nm) | $K_{bond}$ (kJ mol$^{-1}$ nm$^{-2}$) |
|---|---|---|---|
| 18 | 20 | 0.33193 | 40087.22466 |
| 18 | 16 | 0.21976 | 14540.77476 |
| 16 | 1 | 0.34141 | 11331.96785 |
| 2 | 5 | 0.3876 | 20899.64498 |
| 2 | 7 | 0.33386 | 9374.37478 |
| 3 | 6 | 0.37445 | 5658.56784 |
| 6 | 12 | 0.26948 | 8527.95542 |
| 12 | 13 | 0.35298 | 2987.37659 |
| 13 | 14 | 0.42314 | 3968.52702 |
| 14 | 15 | 0.43254 | 3317.70632 |
| 7 | 8 | 0.28111 | 9956.27756 |
| 8 | 9 | 0.34939 | 3021.2073 |
| 9 | 10 | 0.42254 | 3962.00381 |
| 10 | 11 | 0.43329 | 3311.89181 |
| 18 | 21 | 0.36614 | 15885.94134 |
| 21 | 23 | 0.29614 | 5703.90855 |
| 23 | 24 | 0.34514 | 3992.80511 |
| 24 | 25 | 0.36632 | 4720.95029 |
| 25 | 26 | 0.36676 | 4874.236 |
| 26 | 27 | 0.43459 | 3267.08298 |
| 23 | 28 | 0.37121 | 2861.41583 |
| 28 | 29 | 0.41809 | 3880.97892 |
| 29 | 30 | 0.4869 | 2522.95352 |
| 16 | 22 | 0.33178 | 21486.34755 |
| 22 | 31 | 0.32286 | 5666.22972 |
| 31 | 32 | 0.34514 | 4202.04338 |
| 32 | 33 | 0.4207 | 3591.24363 |
| 33 | 34 | 0.48583 | 2478.43986 |
| 31 | 35 | 0.36677 | 2989.03568 |
| 35 | 36 | 0.41871 | 3819.13206 |
| 36 | 37 | 0.48636 | 2522.27919 |
| 17 | 38 | 0.40263 | 31925.8785 |
| 39 | 43 | 0.26255 | 16970.36192 |
| 38 | 42 | 0.29607 | 23851.31594 |
| 40 | 44 | 0.41495 | 8477.0618 |
| 44 | 48 | 0.29313 | 22756.97413 |
| 45 | 49 | 0.26201 | 31206.05154 |

Table S2B Constraint bond parameters

| I | j | length (nm) |
|---|---|---|
| 1 | 2 | 0.36594 |
| 2 | 3 | 0.2937 |
| 1 | 3 | 0.26594 |
| 16 | 17 | 0.37406 |
| 17 | 18 | 0.27251 |
| 16 | 18 | 0.21976 |
| 38 | 39 | 0.33719 |
| 39 | 40 | 0.23918 |
| 38 | 40 | 0.25761 |
| 44 | 45 | 0.33821 |
| 45 | 46 | 0.25799 |
| 46 | 44 | 0.24912 |

Table S2C Flexible bond potential parameters*

*Used for flexible minimization instead of constraints*

| I | j | $b_{eq}$ (nm) | $K_{bond}$ (kJ mol$^{-1}$ nm$^{-2}$) |
|---|---|---|---|
| 1 | 2 | 0.36594 | 5402.99617 |
| 2 | 3 | 0.2937 | 42683.97871 |
| 1 | 3 | 0.26594 | 15894.44095 |
| 16 | 17 | 0.37406 | 16155.77694 |
| 17 | 18 | 0.27251 | 57729.28349 |
| 16 | 18 | 0.21976 | 14540.77476 |
| 38 | 39 | 0.33719 | 46272.12583 |
| 39 | 40 | 0.23918 | 21246.2869 |
| 38 | 40 | 0.25761 | 16615.30225 |
| 44 | 45 | 0.33821 | 45549.01063 |
| 45 | 46 | 0.25799 | 33568.59692 |
| 46 | 44 | 0.24912 | 13902.94084 |

Table S2D Angle potential paramaeters

| I | j | k | $ang_0$ (deg) | $K_{ang}$(kJ $mol^{-1}$ $deg^{-1}$) |
|---|---|---|---|---|
| 23 | 24 | 25 | 134.45984 | 25.36745 |
| 23 | 28 | 29 | 127.94594 | 24.29019 |
| 23 | 25 | 26 | 142.55945 | 22.24468 |
| 25 | 26 | 27 | 143.55522 | 25.89231 |
| 28 | 29 | 30 | 147.4 | 24.89224 |
| 18 | 21 | 23 | 145.99218 | 500 |
| 31 | 32 | 33 | 133.05173 | 21.13936 |
| 31 | 35 | 36 | 129.14664 | 22.67202 |
| 35 | 36 | 37 | 147.37221 | 24.64123 |
| 32 | 33 | 34 | 146.5092 | 23.49165 |
| 16 | 22 | 31 | 132.10041 | 500 |
| 3 | 6 | 12 | 118.8385 | 500 |
| 12 | 13 | 14 | 145.84456 | 28.76617 |
| 13 | 14 | 15 | 148.17859 | 24.91223 |
| 7 | 8 | 9 | 122.7317 | 13.70852 |
| 8 | 9 | 10 | 147.17968 | 28.13906 |
| 9 | 10 | 11 | 148.28177 | 25.08017 |
| 5 | 2 | 1 | 95.52492 | 91.99828 |
| 20 | 18 | 17 | 84.60119 | 442.79611 |
| 17 | 16 | 22 | 162.00306 | 59.24425 |
| 3 | 2 | 7 | 117.64862 | 500 |
| 17 | 16 | 1 | 89.85974 | 218.65071 |
| 18 | 16 | 1 | 133.11893 | 500 |
| 16 | 1 | 3 | 103.54873 | 51.79553 |
| 16 | 1 | 2 | 99.24979 | 45.52129 |
| 40 | 38 | 17 | 95.21793 | 676.88135 |
| 39 | 38 | 17 | 52.93281 | 1061.04971 |
| 38 | 17 | 18 | 161.87875 | 305.40678 |
| 39 | 38 | 17 | 52.93281 | 1061.04971 |
| 38 | 17 | 16 | 130.94667 | 254.45003 |
| 46 | 44 | 40 | 110.89332 | 182.88706 |
| 44 | 40 | 38 | 88.2129 | 386.88439 |
| 45 | 44 | 40 | 64.83742 | 402.35729 |
| 44 | 40 | 39 | 145.59779 | 194.73409 |
| 49 | 45 | 46 | 124.27451 | 286.57197 |
| 48 | 44 | 45 | 89.56246 | 1244.28317 |
| 43 | 39 | 40 | 120.56776 | 148.08985 |
| 42 | 38 | 39 | 88.90197 | 1267.36651 |

Table S2E Dihedral potential paramaeters

| I | j | k | l | $dih_0$ (deg) | $K_{dih}$(kJ $mol^{-1}$ $deg^{-1}$) | multiplicity |
|---|---|---|---|---|---|---|
| 17 | 16 | 1 | 2 | -36.67333 | 12.59321 | 1 |
| 45 | 44 | 40 | 38 | 154.24543 | 7.88236 | 1 |
| 39 | 38 | 17 | 18 | 89.92181 | 6.92723 | 1 |

**Table S3.** Choosing an LJ-modified model of Martini 2 ReLPS such that the difference between the APL of the standard Martini 2 (Head LJ+0) and of the modified Martini 2 model (Head LJ+*n*) is < 5% (n>0).

| **Model** | **Head LJ+0** | **Head LJ+1** | **Head LJ+2** | **Head LJ+3** | **Head LJ+4** | **Head LJ+5** | **Fast** |
|---|---|---|---|---|---|---|---|
| **$A_{LPS}$ ($nm^2$)** | 1.625 ± 0.0115 | 1.648 ± 0.0117 | 1.678 ± 0.0105 | 1.721 ± 0.01 | 1.766 ± 0.00961 | 1.805 ± 0.00942 | 1.774 ± 0.2787 |
| **$\Delta A_{LPS}$ ($nm^2$)** | 0 | 0.023 | 0.053 | 0.096 | 0.141 | 0.18 | 0.149 |
| **$\Delta A_{LPS}$ (%)** | 0 | 1.42 | **3.26** | 5.91 | 8.68 | 11.1 | 9.17 |

**Table S4.** Details on box dimensions per system and GROMACS versions to perform production simulations. Box dimension units are in (X x Y x Z) nm. Note that the box dimensions are shown for 3 replicas at the end of their production simulations, including those systems which were simulated for more than 3 replicas. In the third column, all box dimensions that were setup before energy minimisation are mentioned, except the cells where “eq0” is mentioned in which case box dimensions were obtained after a short equilibration. Abbreviations used in the table are: C36m: CHARMM36m, M2: Martini 2 standard, M2 fast: fast Martini 2, M3: Martini 3, OM: outer membrane.

(page 10-11)

| System | Force field/ LPS type | Box dimension (before minimisation) | Box dimension at end of production replica 1 | Box dimension at end of production replica 2 | Box dimension at end of production replica 3 | Gromacs version used for production simulations |
|---|---|---|---|---|---|---|
| Symmetric ReLPS bilayer | C36m | 15.03662 x 15.03662 x 19.41726 | 14.81480 x 14.81480 x 17.89610 | - | - | 2021.3 |
| Symmetric ReLPS bilayer | M3 | 15.01471x 15.01471 x 25.00000 | 14.69005 x 14.69005 x 15.95020 | 14.65969 x 14.65969 x 16.00532 | 14.67592 x 14.67592 x 15.97799 | 2021.4 |
| Symmetric ReLPS bilayer | M2 | **eq0** 13.75540 13.75540 19.54450 | 13.68184 x 13.68184 x 16.49011 | 13.68082 x 13.68082 x 16.46491 | 13.69249 x 13.69249 x 16.48008 | 2021.4 |
| Symmetric ReLPS bilayer | M2 fast | **eq0** 13.75540 13.75540 19.54450 | 13.68943 x 13.68943 x 16.46326 | 13.68797 x 13.68797 x 16.47290 | 13.69984 x 13.69984 x 16.42741 | 2021.4 |
| OM + DOPC | M2 | 12 x 12 x 16 | 11.72477 x 11.72477 x 15.85321 | 11.72824 x 11.72824 x 15.88695 | 11.75022 x 11.75022 x 15.80153 | 2022.4 |
| OM + DOPC | M2 fast | 12 x 12 x 16 | 11.94381 x 11.94381 x 15.31512 | 11.88679 x 11.88679 x 15.40701 | 11.99261 x 11.99261 x 15.18811 | 2022.4 |
| OM + DOPC | M3 | 12 x 12 x 16 | 12.12834 x 12.12834 x 13.77261 | 12.11392 x 12.11392 x 13.83019 | 12.19810 x 12.19810 x 13.62773 | 2022.4 |
| OM + DPC | M2 | 12 x 12 x 16 | 11.69912 x 11.69912 x 15.90881 | 11.71885 x 11.71885 x 15.90774 | 11.69615 x 11.69615 x 15.93282 | 2022.4 |
| OM + DPC | M2 fast | 12 x 12 x 16 | 11.92728 x 11.92728 x 15.31141 | 11.96242 x 11.96242 x 15.24096 | 11.96738 x 11.96738 x 15.23526 | 2022.4 |
| OM + DPC | M3 | 12 x 12 x 16 | 12.12295 x 12.12295 x 14.00519 | 12.19112 x 12.19112 x 13.85510 | 12.11945 x 12.11945 x 14.04823 | 2022.4 |
| OM + DPC | Steered MD (M2) | 12.15461 x 12.15461 x 14.8195 | 11.98595 x 11.98595 x 15.14625 | 11.99772 x 11.99772 x 15.13335 | 11.97858 x 11.97858 x 15.19924 | 2019.4 |
| OM + DPC | Steered MD (M3) | 12.16435 x 12.16435 x 13.92706 | 12.10633 x 12.10633 x 14.00992 | 12.21149 x 12.21149 x 13.79521 | 12.17416 x 12.17416 x 13.89193 | 2019.4 |
| OM + octane | M2 | 12 x 12 x 16 | 11.58816 x 11.58816 x 16.17199 | 11.72339 x 11.72339 x 15.83037 | 11.63015 x 11.63015 x 16.14074 | 2022.4 |

| | | | | | | |
|---|---|---|---|---|---|---|
| OM + octane | M2 fast | 12 x 12 x 16 | 11.97134 x 11.97134 x 15.27016 | 11.87766 x 11.87766 x 15.44597 | 11.96371 x 11.96371 x 15.25185 | 2021.4 |
| OM + octane | M3 | 12 x 12 x 16 | 12.26741 x 12.26741 x 13.43850 | 12.11925 x 12.11925 x 13.84160 | 12.21953 x 12.21953 x 13.56165 | 2021.4 |
| OmpF + 7 LPS | M2 | 16 x 16 x 14 | 16.53256 x 16.53256 x 12.72750 | 16.66357 x 16.66357 x 12.53920 | 16.68552 x 16.68552 x 12.49366 | 2022.4 |
| OmpF + 7 LPS | M2 fast | 16 x 16 x 14 | 16.69026 x 16.69026 x 12.49359 | 16.71332 x 16.71332 x 12.46103 | 16.70623 x 16.70623 x 12.43382 | 2022.4 |
| OmpF + 7 LPS | M3 | 16 x 16 x 14 | 16.75574 x 16.75574 x 12.00178 | 16.74435 x 16.74435 x 12.02074 | 16.77810 x 16.77810 x 11.95169 | 2022.4 |
| OM + 7x OmpF | M2 | 40 x 40 x 12 | 40.06454 x 40.06454 x 10.59544 | 40.07512 x 40.07512 x 10.63190 | 40.09600 x 40.09600 x 10.53331 | 2020.3 |
| OM + 7x OmpF | M2 fast | 40 x 40 x 12 | 40.78138 x 40.78138 x 10.24050 | 40.87509 x 40.87509 x 10.24242 | 40.83708 x 40.83708 x 10.14357 | 2020.3 |
| OM + 7x OmpF | M2 standard + fast (50/50%) uniform | 40 x 40 x 12 | 38.84201 x 38.84201 x 10.87231 | 38.84872 x 38.84872 x 10.84959 | 38.86150 x 38.86150 x 10.85613 | 2020.3 |
| OM + 7x OmpF | M2 standard + fast (50/50%) non-uniform | 40 x 40 x 12 | 40.10992 x 40.10992 x 10.57044 | 40.00848 x 40.00848 x 10.68509 | 39.96832 x 39.96832 x 10.60062 | 2020.3 |
| OMP island | M2 | 150 x 150 x 30 | 146.07164 x 146.07164 x 27.57932 | 146.80037 x 146.80037 x 27.30904 | 146.83083 x 146.83083 x 27.29945 | 2022.4 |
| OMP island | M2 fast | 150 x 150 x 30 | 150.12053 x 150.12053 x 26.10580 | 149.97099 x 149.97099 x 26.15880 | 150.06816 x 150.06816 x 26.13442 | 2022.4 |
| OMP island | M3 | 150 x 150 x 30 | 141.61624 x 141.61624 x 30.00586 | 140.52150 x 140.52150 x 30.47952 | 142.38239 x 142.38239 x 29.68060 | 2022.4 |
| OM + PMB1 | M2 | 20 x 20 x 12 | 20.2 x 20.2 x 11.8 | 20.2 x 20.2 x 11.8 | 20.2 x 20.2 x 11.8 | 2021.5 |
| OM + PMB1 | M2 fast | 20 x 20 x 12 | 20.8 x 20.8 x 11.1 | 20.6 x 20.6 x 11.4 | 20.5 x 20.5 x 11.4 | 2021.5 |

#### 3. Supplementary Figures

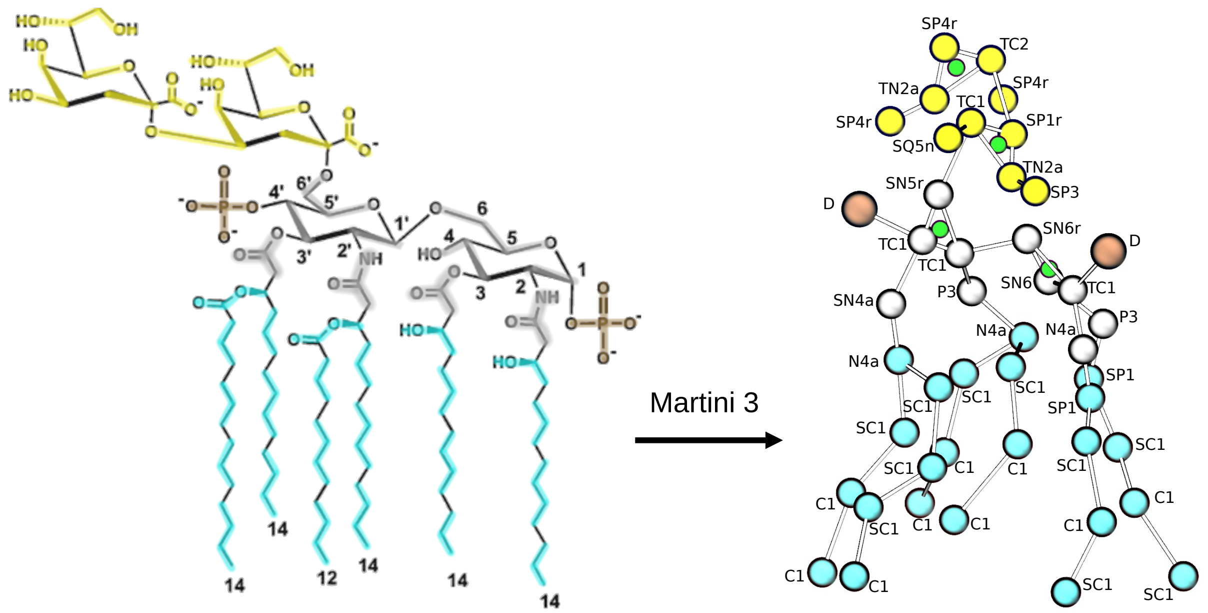

**S1. Martini 3 ReLPS mapping and associated bead types.** Groups of two to four heavy atoms were mapped into coarse-grained (CG) beads. Notably the sugar rings were modelled with an extra virtual site bead (depicted in green) to better characterise their volume and interactions. Here, each highlighted segment of the 2D depiction corresponds to the same-coloured bead on the right CG molecule taken from a simulation snapshot. The four different colours represent: Kdo sugars i.e. first two in core region (yellow), GlcN sugars from lipid A moiety (grey), phosphate substituents (brown), acyl chains (cyan). The labels on the CG representation show the final assigned beadtype.

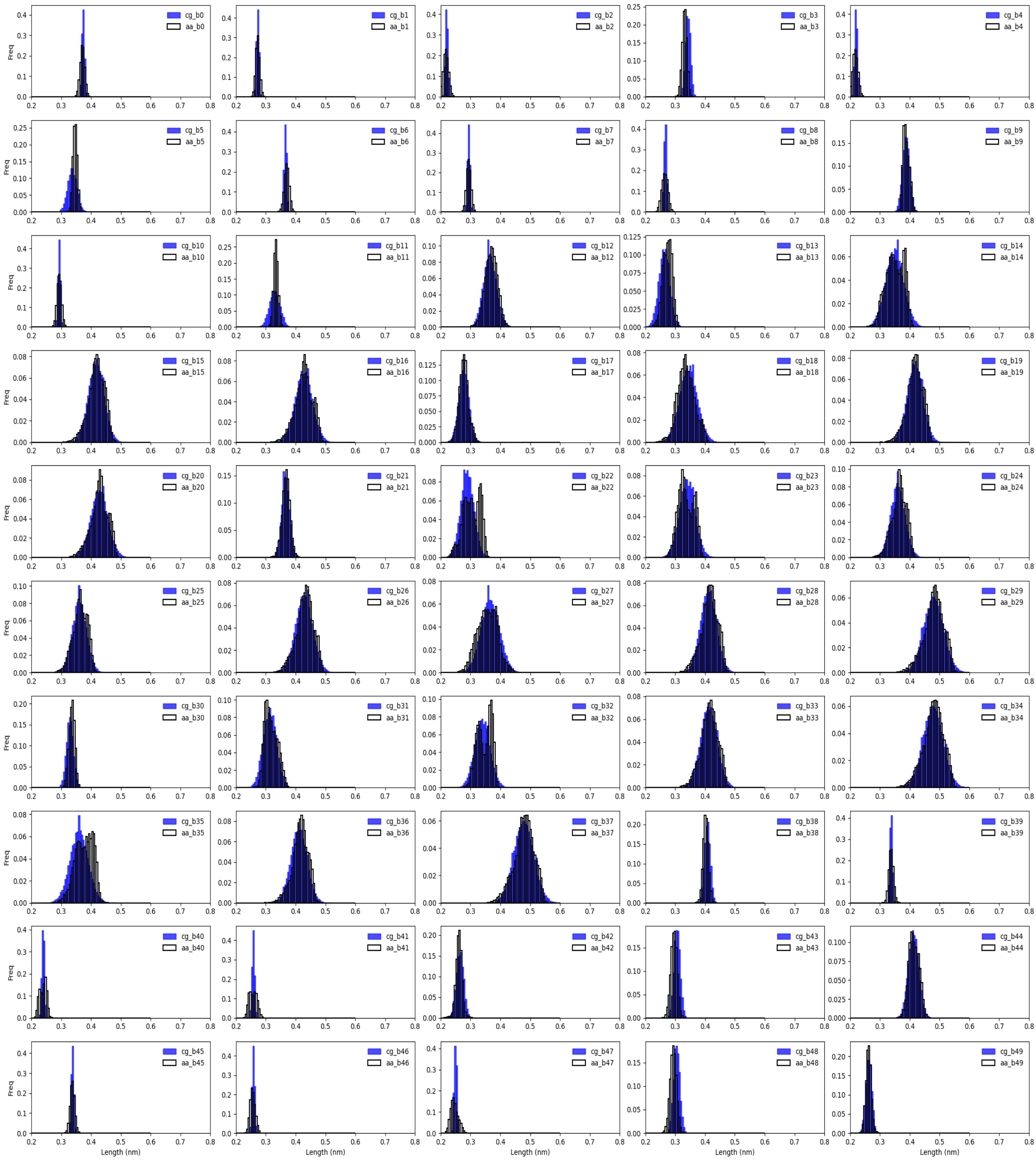

**S2. Martini 3 ReLPS bond distributions**. Distributions for all the bonds are plotted separately. Blue bins correspond to the Martini 3 model whereas black outlines correspond to the data obtained from pseudo-coarse-grained atomistic simulation (i.e converting the atomistic coordinates into coarse-grain model for each of the frames).

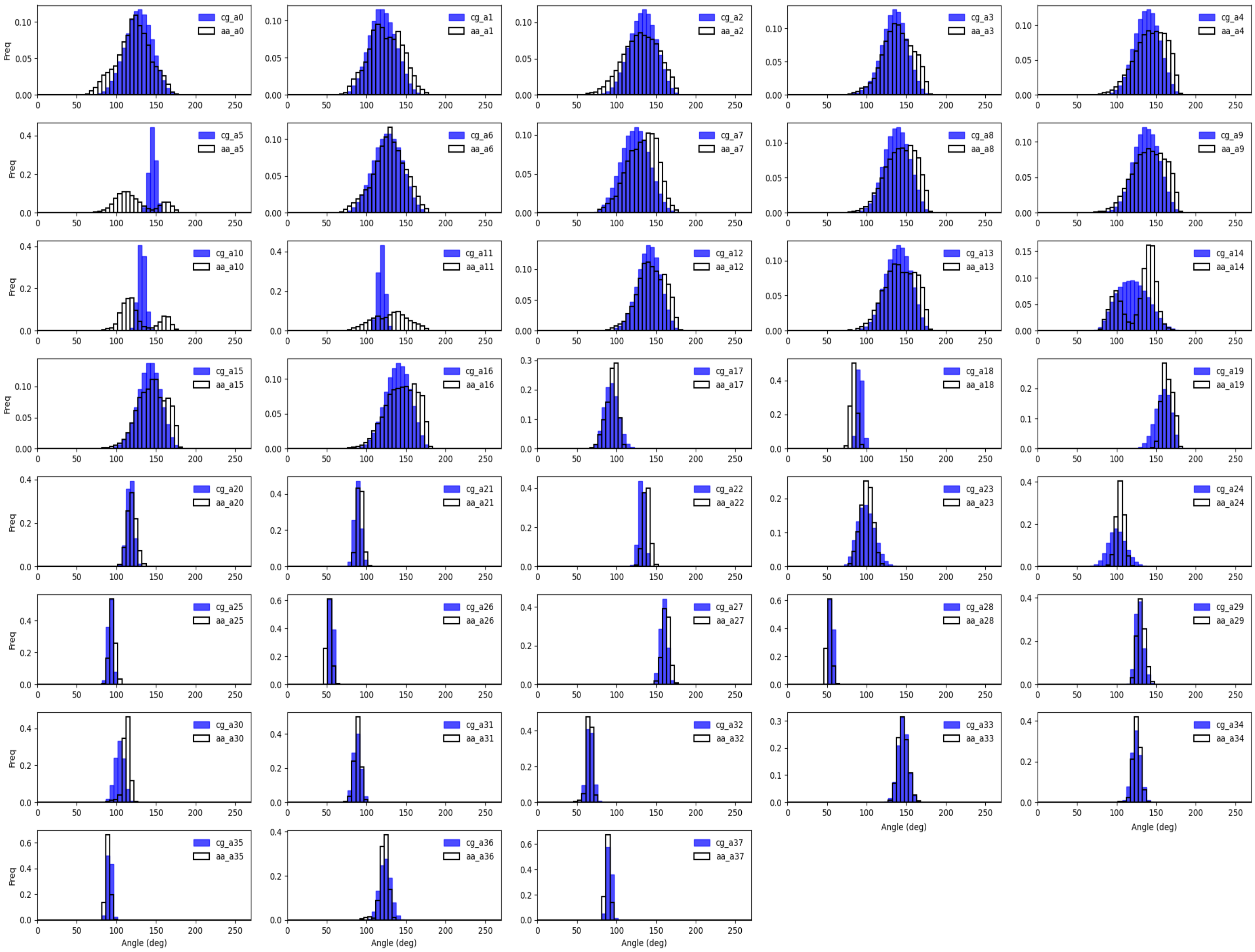

**S3. Martini 3 ReLPS angle distributions**. Distributions for all the angles are plotted separately. Blue bins correspond to the Martini 3 model whereas black outlines correspond to the data obtained from pseudo-coarse-grained atomistic simulation (i.e converting the atomistic coordinates into coarse-grain model for each of the frames).

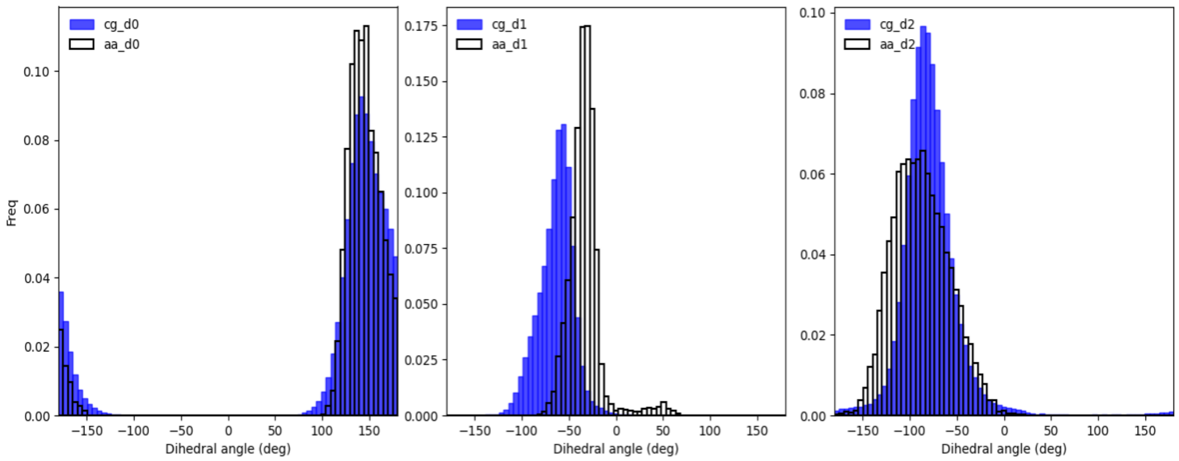

**S4. Martini 3 ReLPS dihedral distributions.** Distributions for all the angles are plotted separately. Blue bins correspond to the Martini 3 model whereas black outlines correspond to the data obtained from pseudo-coarse-grained atomistic simulation (i.e. atomistic coordinates converted into coarse-grained models for each of the frames).

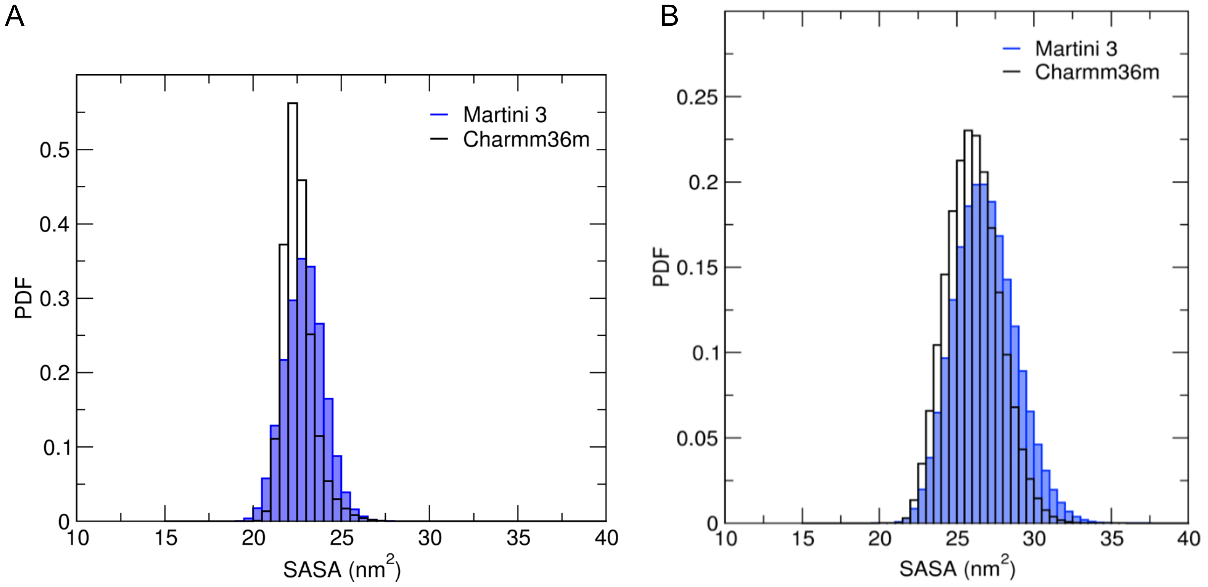

**S5. ReLPS Solvent-accessible surface area (SASA) distribution.** Probability distribution functions (PDF) of the SASA of a single ReLPS **(A)** in solution and **(B)** in a symmetric bilayer are plotted in blue and compared with their atomistic counterpart in black (CHARMM36m).

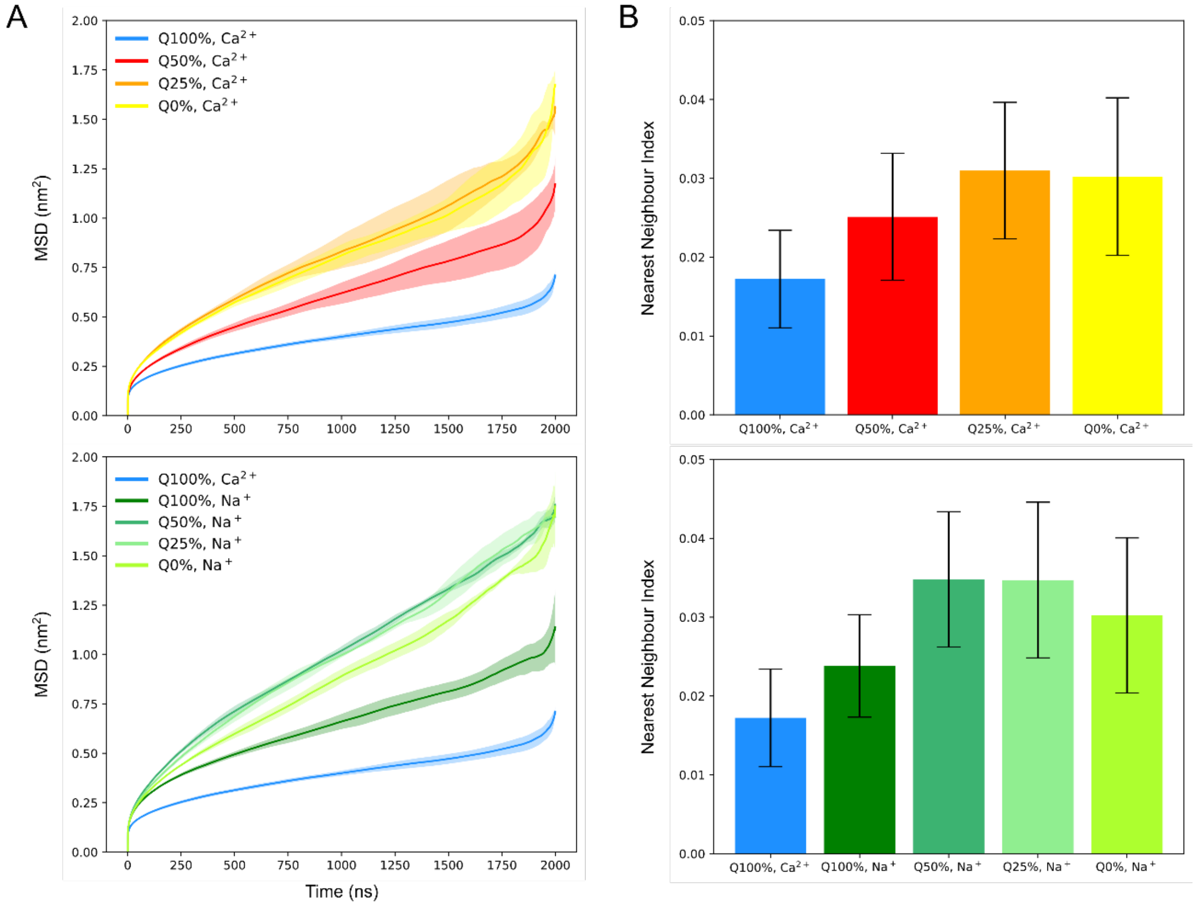

**S6. Evaluating the influence of scaling down charges on Martini 2 ReLPS beads and cations on its (A) MSD and (B) NNI.** For each plot, the data shown are calculated from three simulation replicas. In (A), the dark line represents the mean value from the three trajectories and the shaded regions indicate the standard deviation. Systems are labelled according to the modifications made to the anionic charges of ReLPS beads and the ions used to neutralise the system. For example, “Q50%, $Na^{+}$” means that the negative charges on all anionic ReLPS beads were scaled down to 50% and the system was neutralised by monovalent cations in addition to 0.15 M NaCl. The MSD and NNI data are split into two plots each for visual clarity.

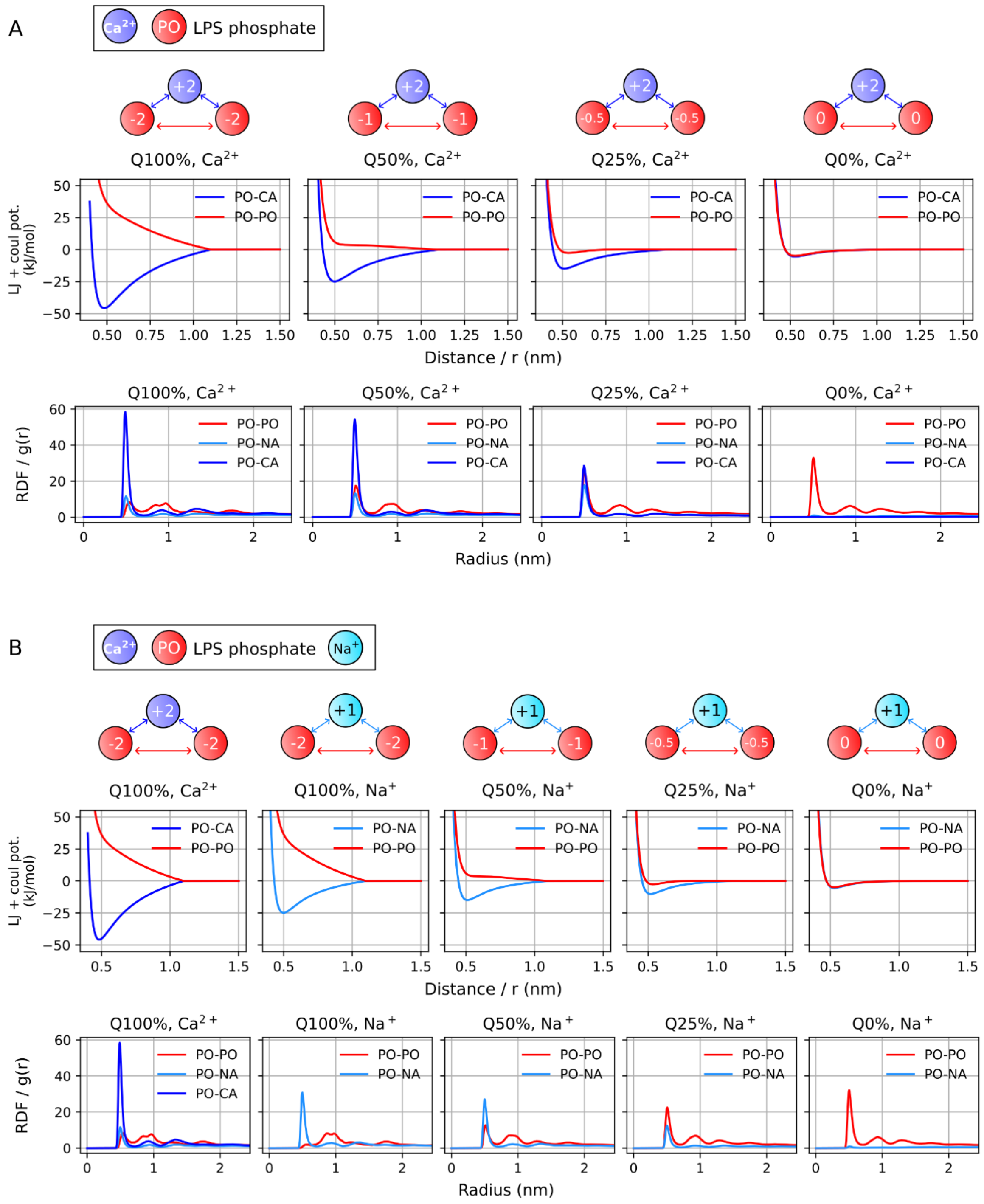

**S7. Reasoning for MSD and NNI of ReLPS approaching a peak and then decreasing when their phosphate bead charges are scaled from 100% to 50%, 25% and 0%.** Total potential energies (LJ + coulombic potentials) (top row) and RDFs (bottom row) of monovalent, divalent cations, and other LPS phosphates (PO beads) with reference to LPS phosphates, labelled as 'PO-NA', 'PO-CA', 'PO-PO' respectively. The system charge neutralising cations are **(A)** the divalent cations ($Ca^{2+}$), and **(B)** the monovalent cations ($Na^{+}$).

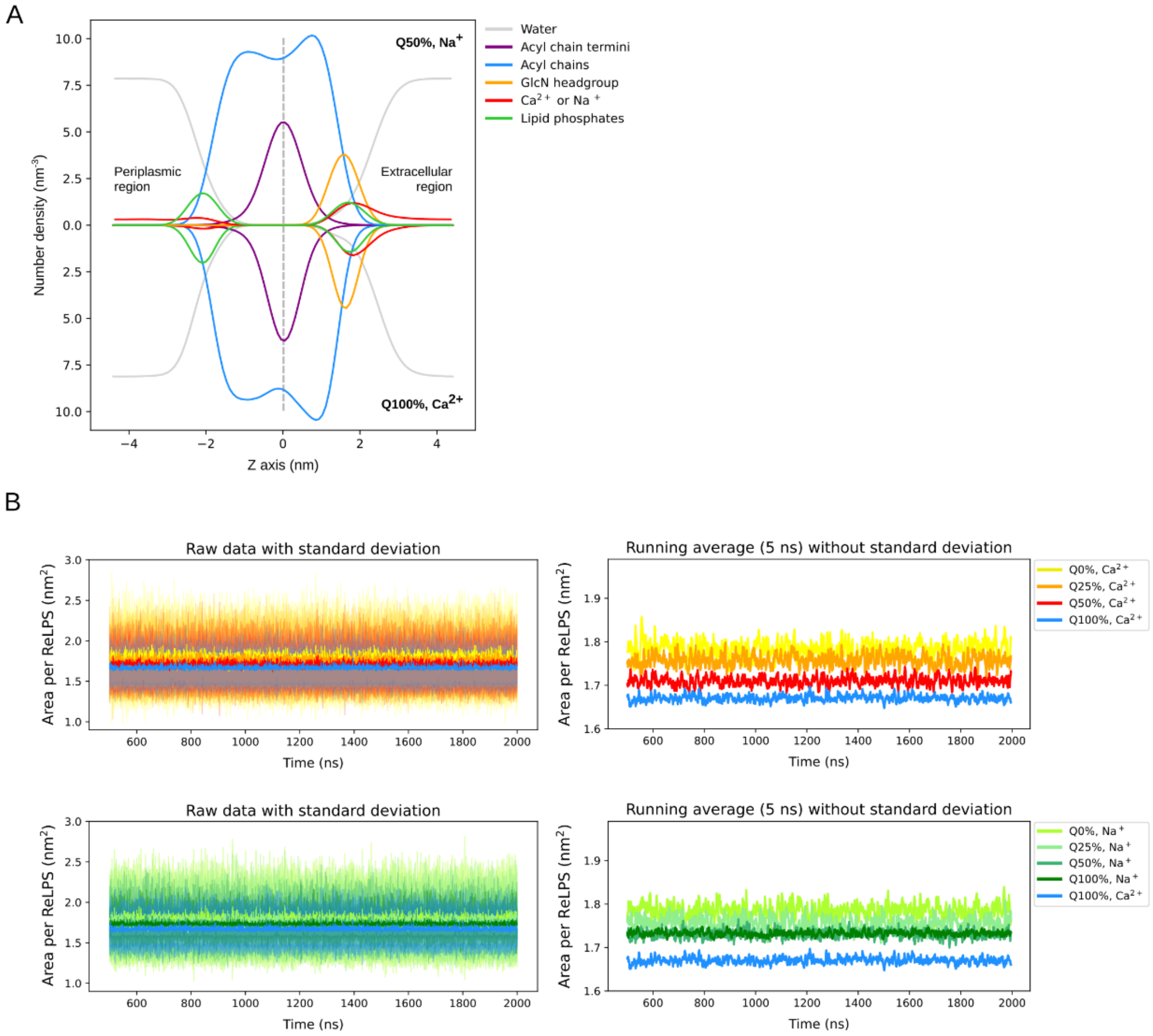

**S8. Evaluating the influence of only scaling down charges on Martini 2 ReLPS beads and cations on its partial density profiles and area per lipid. (A)** Partial density profiles of various components of the 'Q50%, $Na^{+}$' system (top half) compared to the (unmodified) reference system 'Q100%, $Ca^{2+}$' (bottom half). The density profiles are such that the hydrophobic core of the membrane is located at the i.e. Z distance = 0 nm (grey dotted line), and the periplasmic and extracellular regions are located on the left and right hand side of the dotted line respectively. **(B)** APL versus time averaged across simulation replicas of the various systems. The reference system 'Q100%, $Ca^{2+}$' is shown in blue. For visual clarity, in addition to the APL with standard deviation (left), we show the running average (over 5 ns windows) without standard deviation (right). The plots are further split into two showing APL data from simulations with $Ca^{2+}$ and $Na^{+}$ neutralising cations.

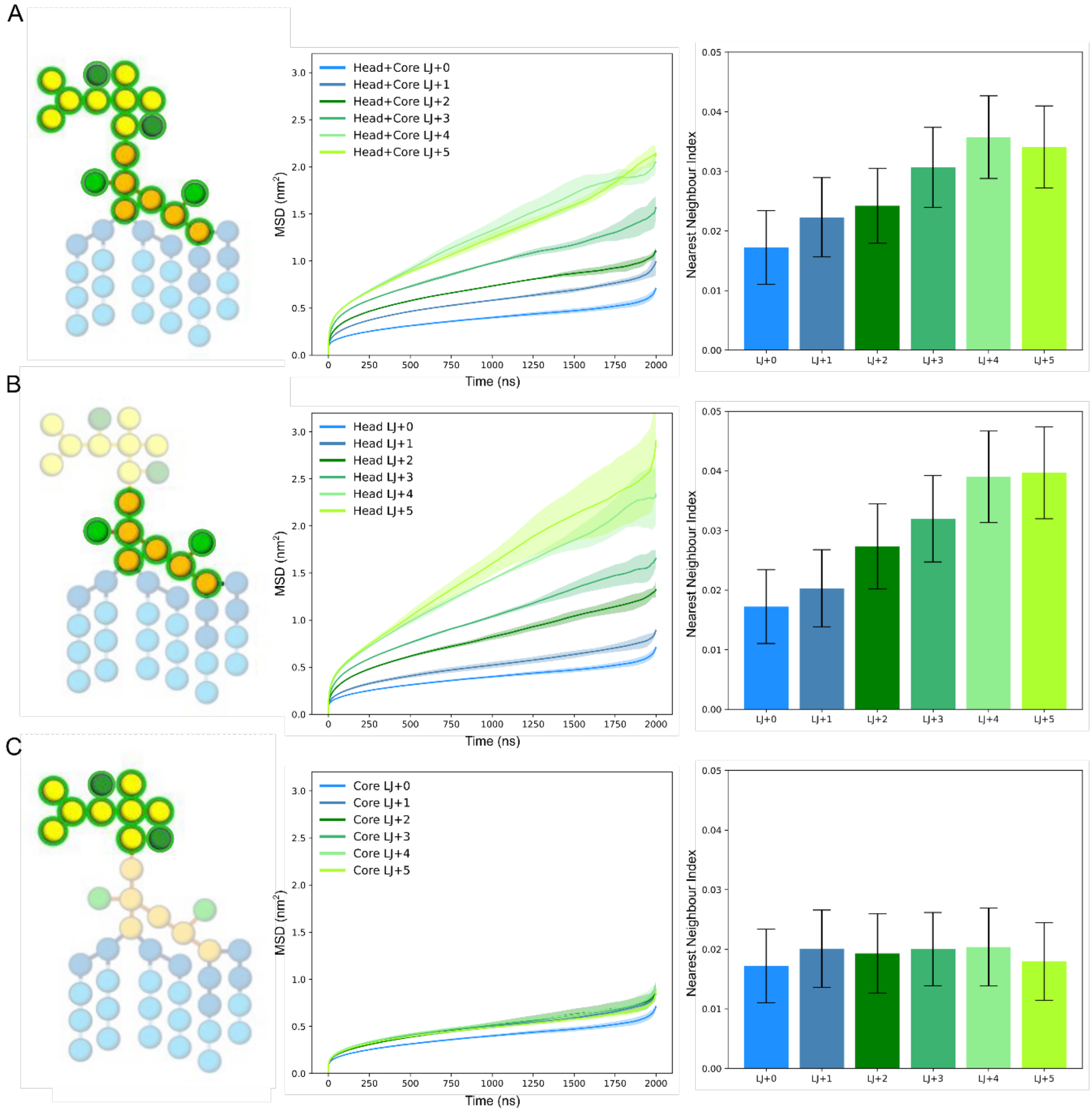

**S9. Evaluating the influence of only scaling down LJ potentials on Martini 2 ReLPS beads on its MSD and NNI.** MSD and NNI data are shown in the middle and on the right respectively for each combination of LJ-modified beads (scaled down up to 5 levels) in the Martini 2 ReLPS molecule i.e. **(A)** the GlcN headgroup and the Kdo core, **(B)** only the GlcN headgroup, **(C)** only the Kdo core. The LJ-modified beads are highlighted and shown within a green circle in the schematic diagrams on the left.

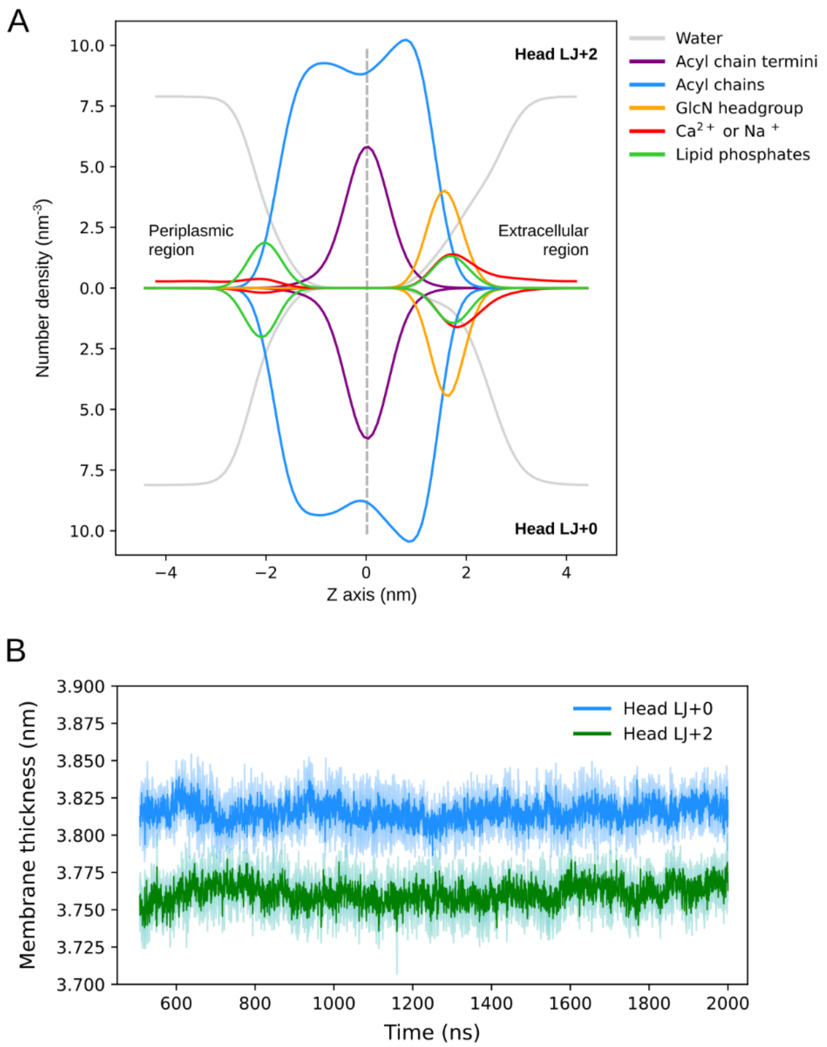

**S10. Evaluating the influence of scaling down modifying LJ potentials on Martini 2 ReLPS beads on its partial density profiles and membrane thickness. (A)** Partial density profiles of various components of the 'Head LJ+2' system (top half) compared to those of the (unmodified) reference system 'Q100%, $Ca^{2+}$' (bottom half). The density profiles are centered such that the hydrophobic core of the membrane is located at the center i.e. Z distance = 0 nm (grey dotted line), and the periplasmic and extracellular regions are located on the left and right hand side of the center respectively. **(B)** Membrane thickness versus time (starting from 500 ns) calculated for the 'Head LJ+2' system (dark green) and compared to that of the reference system 'Head LJ+0' (blue).

A

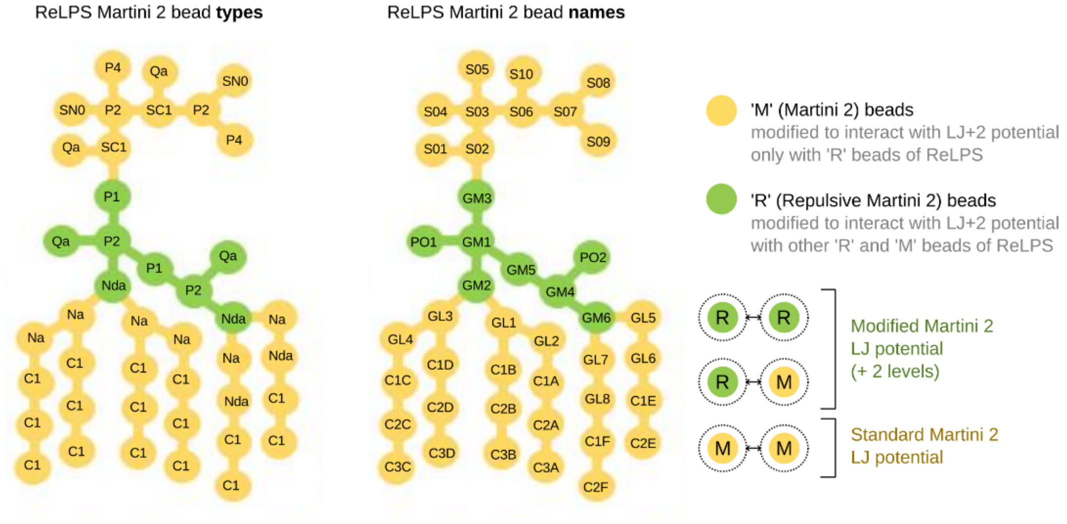

B

**Standard Martini 2 ReLPS**

**Fast Martini 2 ReLPS**

R ⟷ R LJ potential level

| | Qa | Nda | P1 | P2 |
|---|---|---|---|---|
| Qa | 1.0 | 1.0 | 1.0 | 1.0 |
| Nda | 1.0 | 2.0 | 2.0 | 2.0 |
| P1 | 1.0 | 2.0 | 2.0 | 2.0 |
| P2 | 1.0 | 2.0 | 2.0 | 2.0 |

| | RQa | RNda | RP1 | RP2 |
|---|---|---|---|---|
| RQa | 3.0 | 3.0 | 3.0 | 3.0 |
| RNda | 3.0 | 4.0 | 4.0 | 4.0 |
| RP1 | 3.0 | 4.0 | 4.0 | 4.0 |
| RP2 | 3.0 | 4.0 | 4.0 | 4.0 |

M ⟷ R LJ potential level

| | Qa | Nda | P1 | P2 |
|---|---|---|---|---|
| Qa | 1.0 | 1.0 | 1.0 | 1.0 |
| Na | 3.0 | 2.0 | 2.0 | 2.0 |
| Nda | 1.0 | 2.0 | 2.0 | 2.0 |
| P2 | 1.0 | 2.0 | 2.0 | 2.0 |
| P4 | 0.0 | 3.0 | 2.0 | 2.0 |
| C1 | 9.0 | 6.0 | 6.0 | 7.0 |
| SC1 | 9.0 | 6.0 | 6.0 | 7.0 |
| SN0 | 4.0 | 4.0 | 3.0 | 3.0 |

| | RQa | RNda | RP1 | RP2 |
|---|---|---|---|---|
| MQa | 3.0 | 3.0 | 3.0 | 3.0 |
| MNa | 5.0 | 4.0 | 4.0 | 4.0 |
| MNda | 3.0 | 4.0 | 4.0 | 4.0 |
| MP2 | 3.0 | 4.0 | 4.0 | 4.0 |
| MP4 | 2.0 | 5.0 | 4.0 | 4.0 |
| MC1 | 9.0 | 8.0 | 8.0 | 9.0 |
| MSC1 | 9.0 | 8.0 | 8.0 | 9.0 |
| MSN0 | 6.0 | 6.0 | 5.0 | 5.0 |

**LJ potential levels defined in the standard Martini 2 FF**

| | |
|---|---|
| 0.0 | supra attractive |
| 0.5 | |
| 1.0 | attractive |
| 1.5 | |
| 2.0 | almost attractive |
| 2.5 | |
| 3.0 | semi attractive |
| 3.5 | |
| 4.0 | intermediate |
| 4.5 | |
| 5.0 | almost intermediate |
| 5.5 | |
| 6.0 | semi repulsive |
| 6.5 | |
| 7.0 | almost repulsive |
| 7.5 | |
| 8.0 | repulsive |
| 8.5 | |
| 9.0 | super repulsive |

**S11. Schematic illustrations showing the modified Martini 2 bead names and types, and the Lennard-Jones (LJ) interaction matrix compared between the standard and fast Martini 2 ReLPS parameters. (A)** ReLPS labelled as per the standard Martini 2 bead names and types. These beads are coloured according to our parametrisation of the fast Martini 2 ReLPS. The headgroup beads (green) are named with a prefix 'R' given that they undergo a repulsion of 2 levels compared to standard Martini 2, and all other beads are named 'M' given that they had to be modified considering their potential interactions with the 'R' beads. **(B)** Comparison of LJ interaction matrices between the standard and fast Martini 2 ReLPS beads. LJ levels 0 to 9 are coloured on a blue-white-red (low-high) scale.

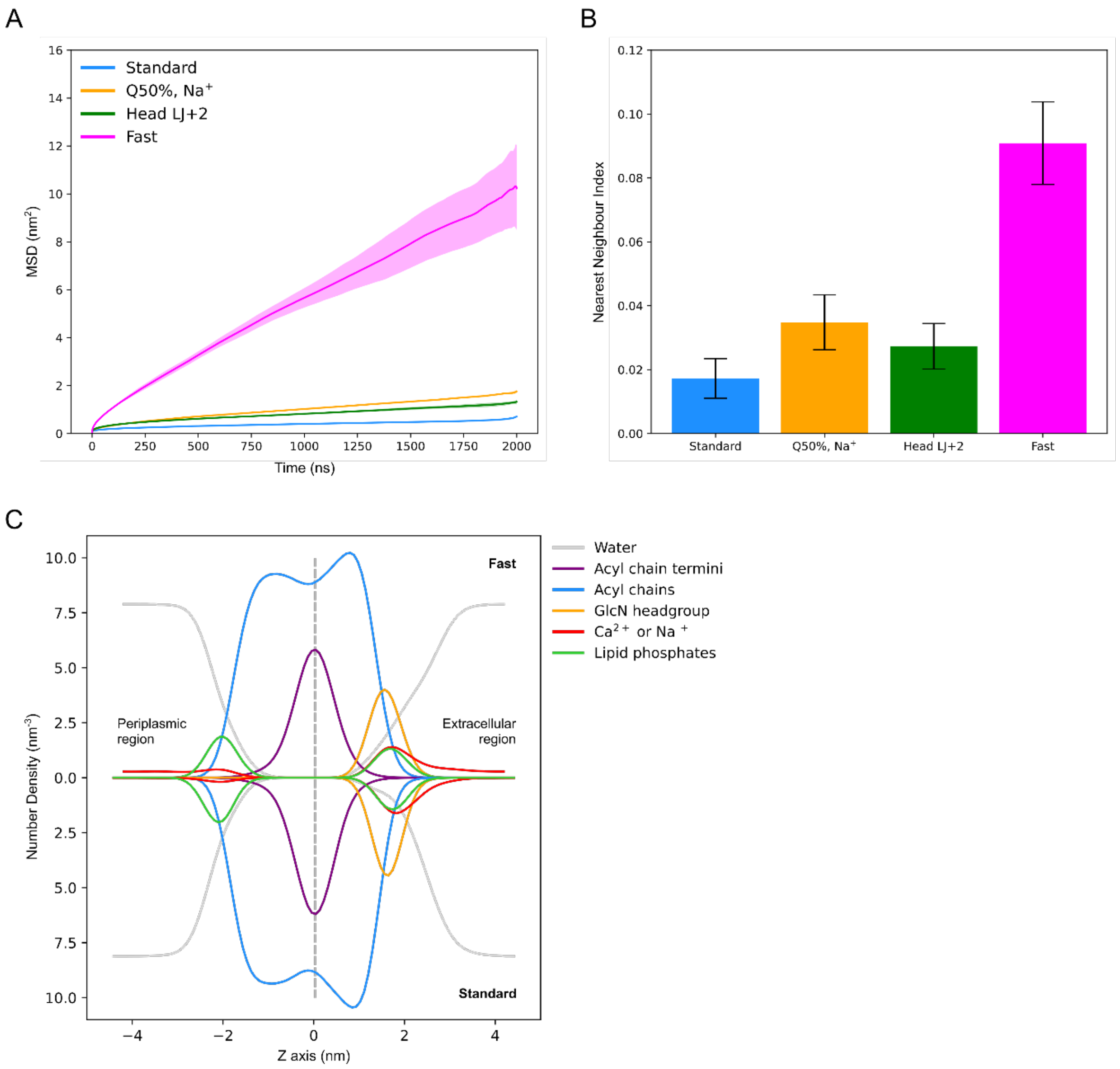

**S12. Evaluating the influence of the fast Martini 2 ReLPS parameters on its MSD, NNI, and partial densities. (A)** Comparing the MSDs and **(B)** NNI of (1) fast Martini 2 ReLPS (magenta), (2) ReLPS with only their charges modified i.e. ‘Q50% $Na^{+}$’ (orange), (3) ReLPS with only their LJ modified i.e. ‘Head LJ+2’ (green), and (4) the reference system with standard Martini 2 parameters (blue). **(C)** Partial density profiles of various components of the system containing fast Martini 2 ReLPS (top half of the plot) compared to those of the standard Martini 2 (unmodified) system (bottom half of the plot). The density profiles are such that the hydrophobic core of the membrane is located at the i.e. Z distance = 0 nm (grey dotted line), and the periplasmic and extracellular regions are located on the left and right hand side of the centre respectively.

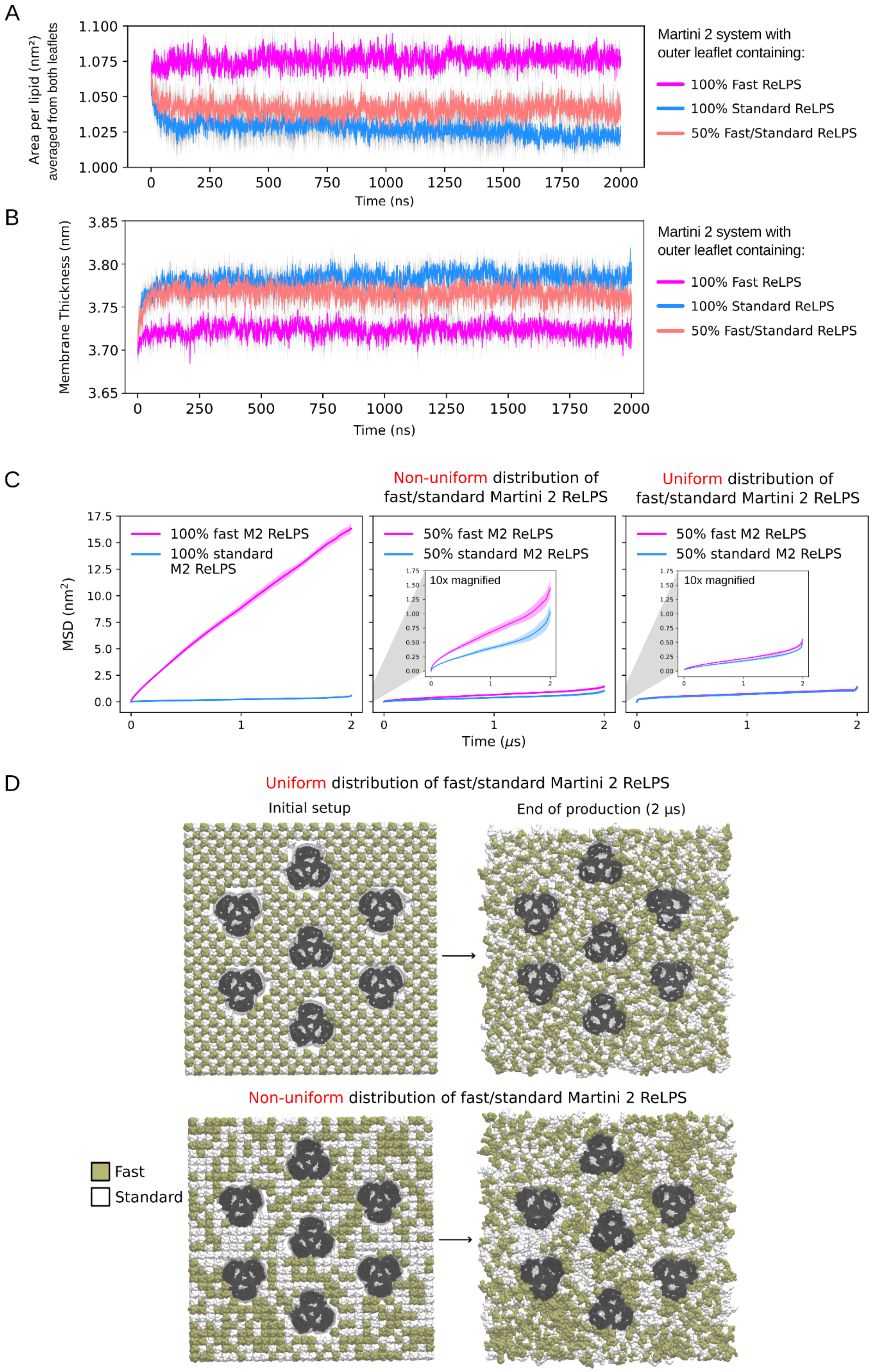
A
Area per lipid (nm²) averaged from both leaflets
Time (ns)
Martini 2 system with outer leaflet containing:
100% Fast ReLPS
100% Standard ReLPS
50% Fast/Standard ReLPS
B
Membrane Thickness (nm)
Time (ns)
Martini 2 system with outer leaflet containing:
100% Fast ReLPS
100% Standard ReLPS
50% Fast/Standard ReLPS
C
Non-uniform distribution of fast/standard Martini 2 ReLPS
Uniform distribution of fast/standard Martini 2 ReLPS
MSD (nm²)
100% fast M2 ReLPS
100% standard M2 ReLPS
50% fast M2 ReLPS
50% standard M2 ReLPS
10x magnified
Time (μs)
D
Uniform distribution of fast/standard Martini 2 ReLPS
Initial setup
End of production (2 μs)
Non-uniform distribution of fast/standard Martini 2 ReLPS
Fast
Standard

**S13. Comparing APL, membrane thickness, and MSDs of Martini 2 systems containing OMPs surrounded by 100% fast ReLPS, 100% standard ReLPS, and 50% fast/standard ReLPS in the outer leaflet. (A)** Averaged APL from both membrane leaflets and **(B)** membrane thickness versus time of systems containing 100% fast ReLPS (magenta), 100% standard ReLPS (cyan), and 50% fast/standard ReLPS (light red) in the outer leaflet of the outer membrane model. Standard deviations for each data line is shown in light grey. **(C)** MSDs versus time compared between 100% fast and 100% standard Martini 2 (left). In the 50% fast/standard ReLPS-containing systems, two initial setups were obtained i.e. a non-uniform and a uniform distribution of Martini 2 fast/standard ReLPS. From each of these simulations, MSDs were calculated separately for the fast ReLPS molecules and for the standard ReLPS molecules. A comparison of MSDs between the fast and standard ReLPS molecules is made from the simulations starting with the non-uniform distribution (center) and the uniform distribution (right). **(D)** Snapshots of the initial setup (left) and final frame (right) of simulations starting with a uniform distribution (top) and non-uniform distribution (bottom) of fast and standard ReLPS molecules.

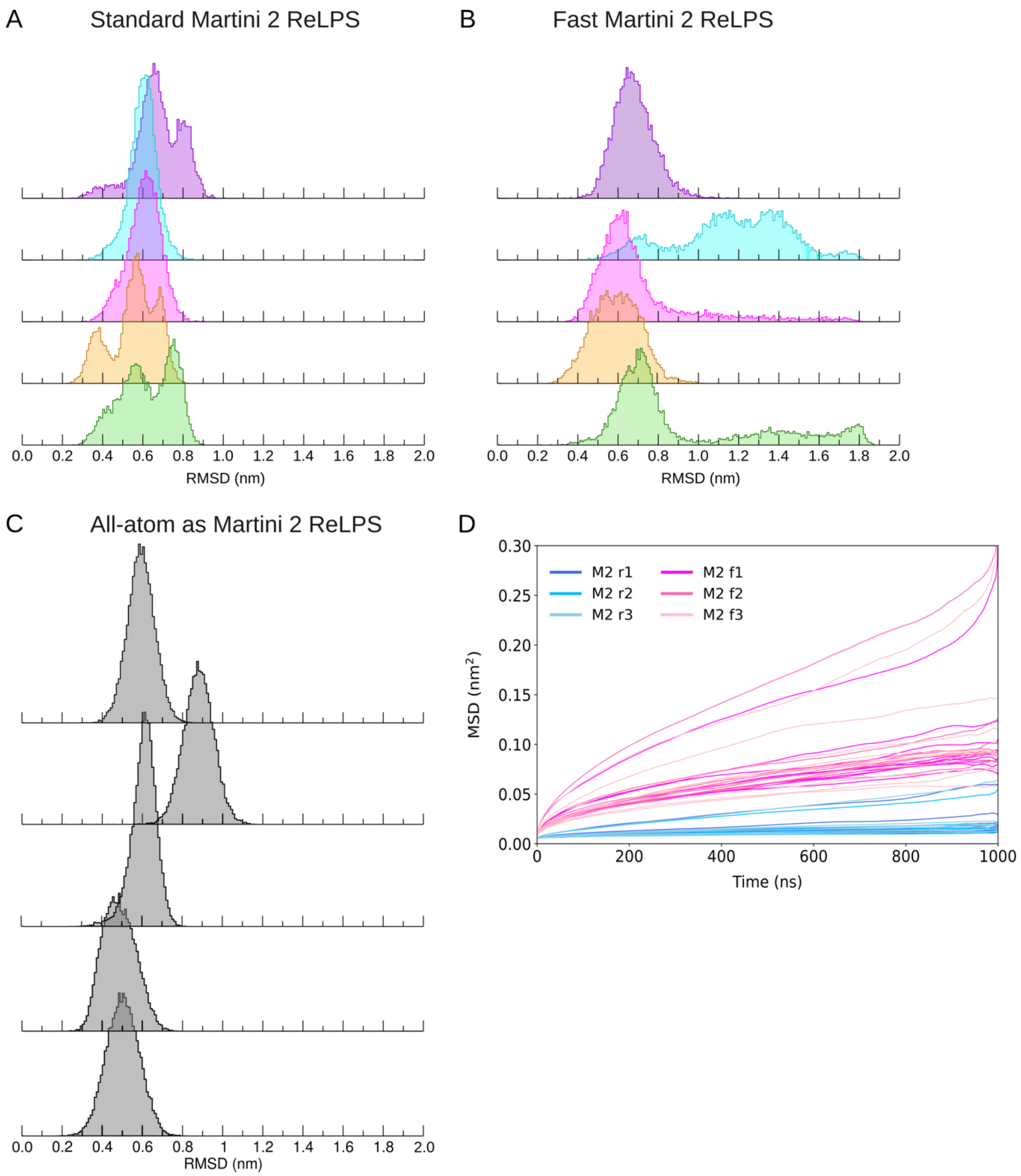

**S14. Internal flexibility and lateral mobility of ReLPS in a 100% ReLPS symmetric bilayer. (A,B,C)** RMSD histograms for 5 individual ReLPS molecules. The full 10 μs trajectory was analysed for the **(A)** standard Martini 2 and **(B)** fast Martini 2 models. RMSD was calculated on the whole molecule after fitting in the central region (GlcN and phosphates). The atomistic simulation was mapped into Martini 2 where 5 randomly chosen individual ReLPS molecules were analysed as explained and shown in **(C)**. **(D)** Lateral MSD of ReLPS in replicas of standard (M2 r1-3) and fast Martini 2 (M2 f1-3) ReLPS computed independently for 10 non-overlapping 1 μs-long trajectory segments.

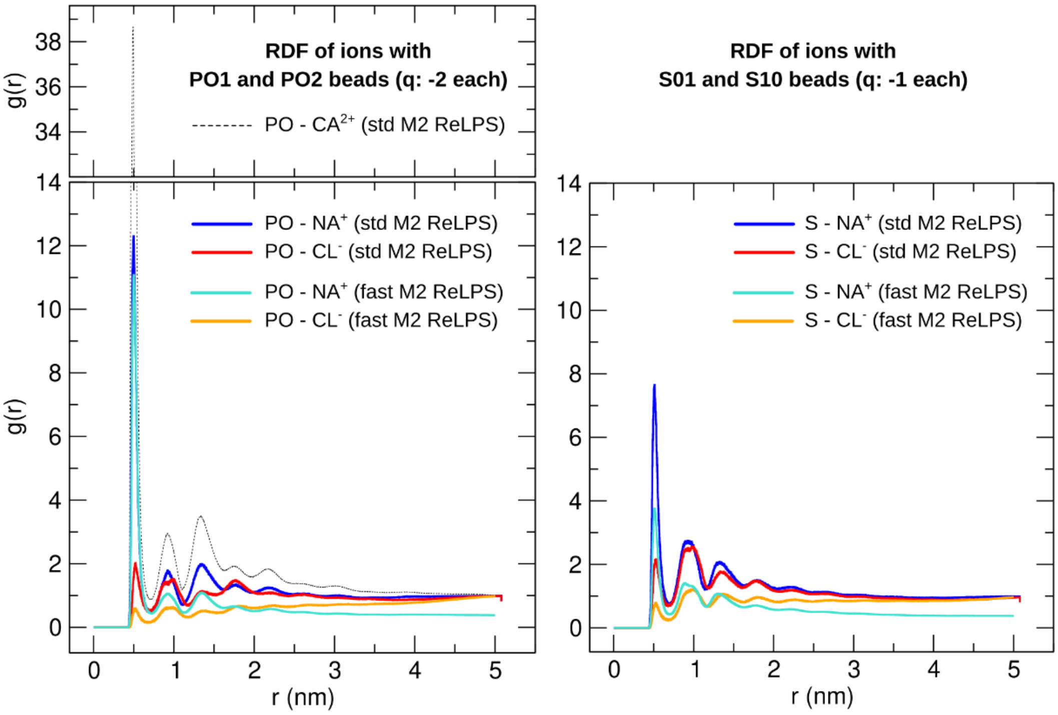

**S15. Comparing ReLPS phosphate-ion interactions between standard and fast Martini 2 models.** RDFs of (1) divalent, monovalent cations, and monovalent anions with ReLPS phosphate beads (left), and (2) monovalent cations and anions with charged glycan (S) beads in the ReLPS core.

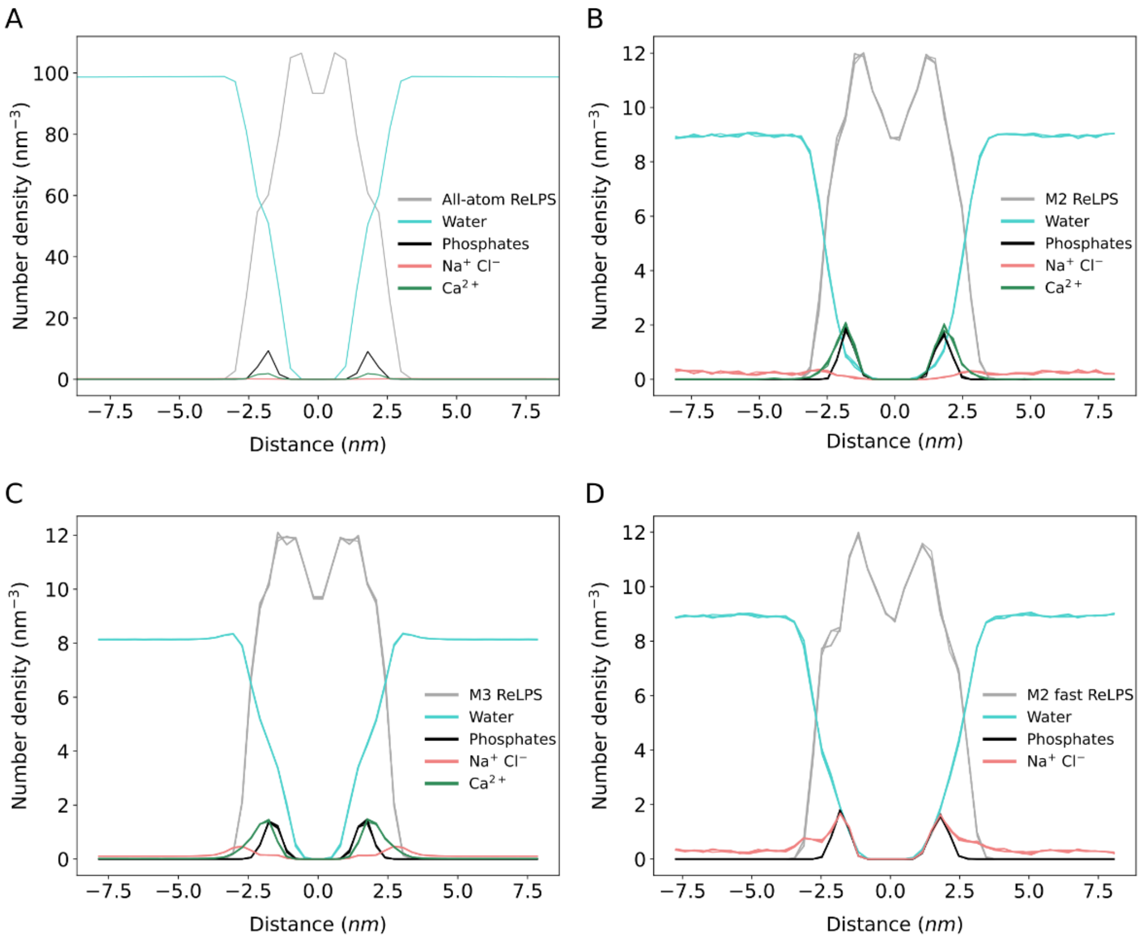

**S16. Lateral molecular organisation of a 100% ReLPS symmetric bilayer.** Lateral number densities of different molecular groups of interest along the membrane normal, computed for **(A)** the all-atom simulation, **(B)** standard Martini 2**, (C)** Martini 3, and **(D)** fast Martini 2. Distance = 0 nm corresponds to the centre of the bilayer along the Z axis.

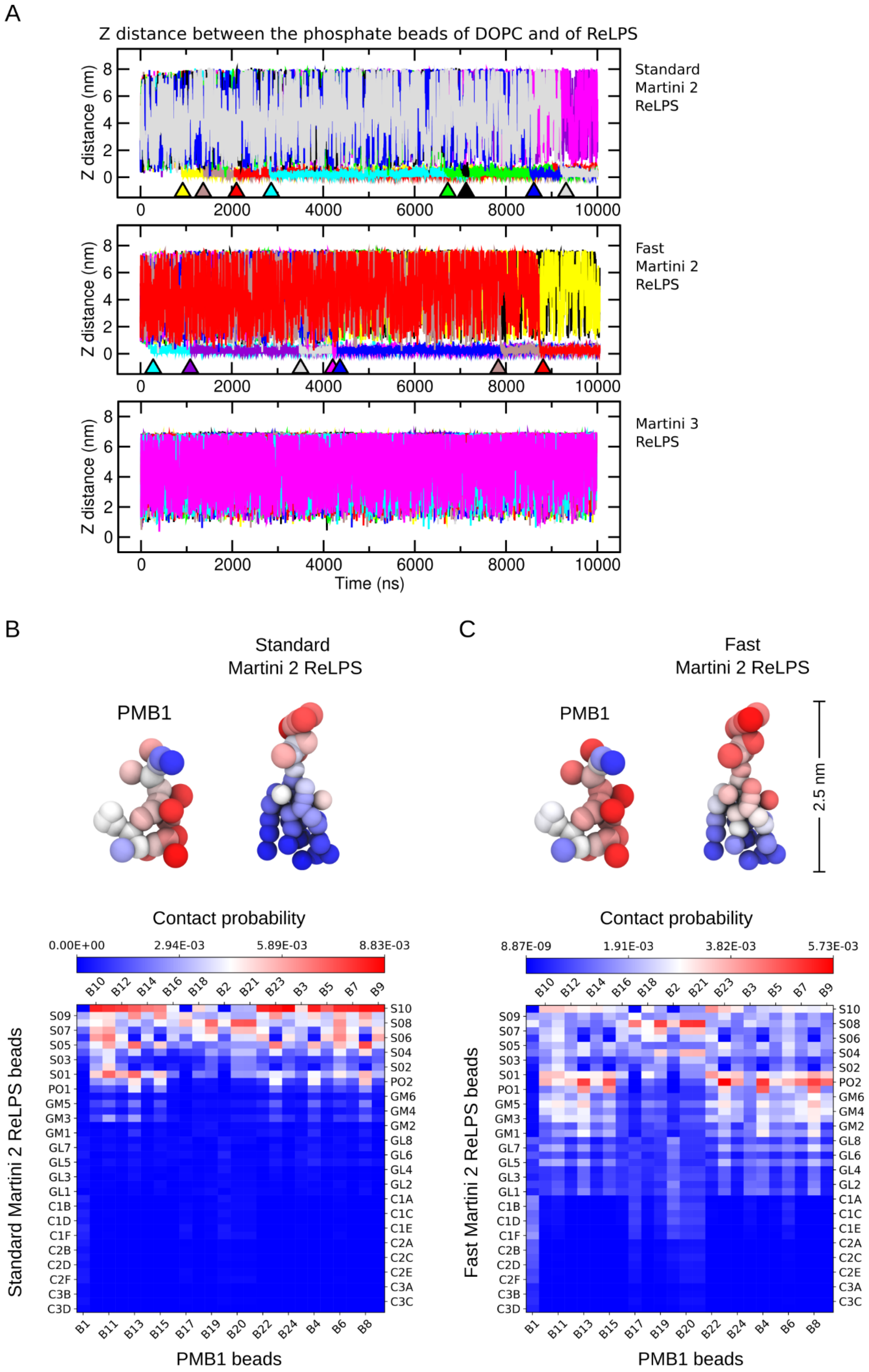

A
Z distance between the phosphate beads of DOPC and of ReLPS
Z distance (nm)
Standard Martini 2 ReLPS
Fast Martini 2 ReLPS
Martini 3 ReLPS
Time (ns)
B
C
Standard Martini 2 ReLPS
Fast Martini 2 ReLPS
PMB1
2.5 nm
Contact probability
0.00E+00
2.94E-03
5.89E-03
8.83E-03
8.87E-09
1.91E-03
3.82E-03
5.73E-03
Standard Martini 2 ReLPS beads
Fast Martini 2 ReLPS beads
PMB1 beads

**S17. DOPC and PMB1 insertion tests conducted with standard and fast Martini 2 ReLPS. (A)** Distance along the Z dimension (perpendicular to the membrane) versus time calculated between the phosphate bead(s) of the (single) DOPC molecule and of the ReLPS molecules in the outer leaflet of the membrane. **(B)** Average number of contacts of PMB1 beads with standard Martini 2 ReLPS beads are mapped onto their respective coarse-grained structures on a blue-white-red (low-high) scale. **(C)** The same is performed for contacts between PMB1 and fast Martini 2 ReLPS. For both (A) and (B), their respective contact probabilities are plotted on a matrix and shown below using the same colour scale. For visual clarity, labels are alternated between opposite axes.

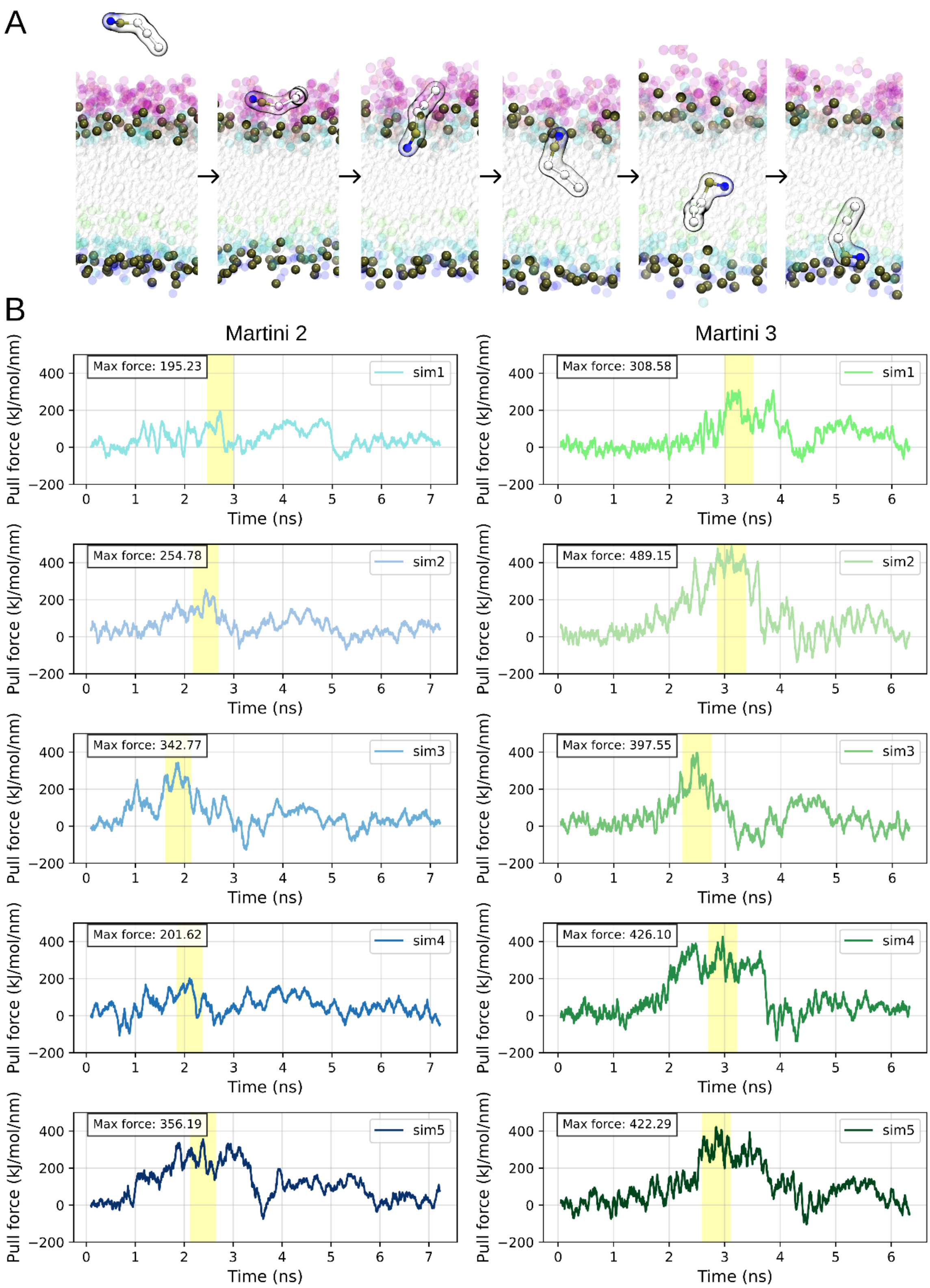

**S18. Steered MD simulations of DPC. (A)** Snapshots taken from steered MD simulations of DPC in Martini 3. **(B)** Data showing the change in pulling force versus time while DPC was steered from water into the membrane using Martini 2 (left) and Martini 3 ReLPS (right) models. In each plot, the maximum force experienced by DPC is labelled and corresponds to the yellow region in each plot.

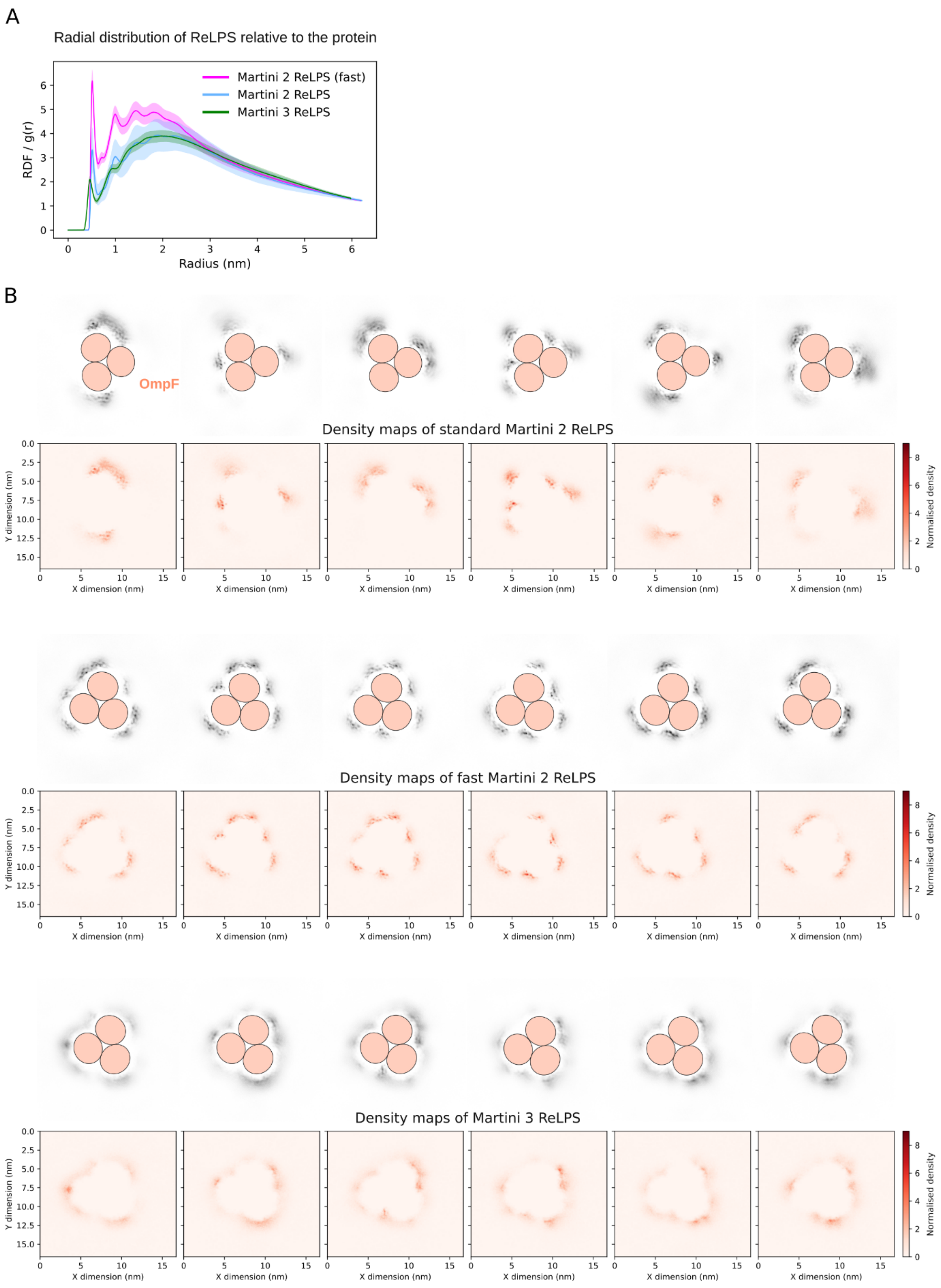

**S19. Densities of ReLPS relative to OmpF in an OM model with 2% ReLPS in the outer leaflet.** (A) Radial distribution functions and (B) 2D density maps of the Martini 2 and 3 ReLPS models with reference to an OmpF trimer.

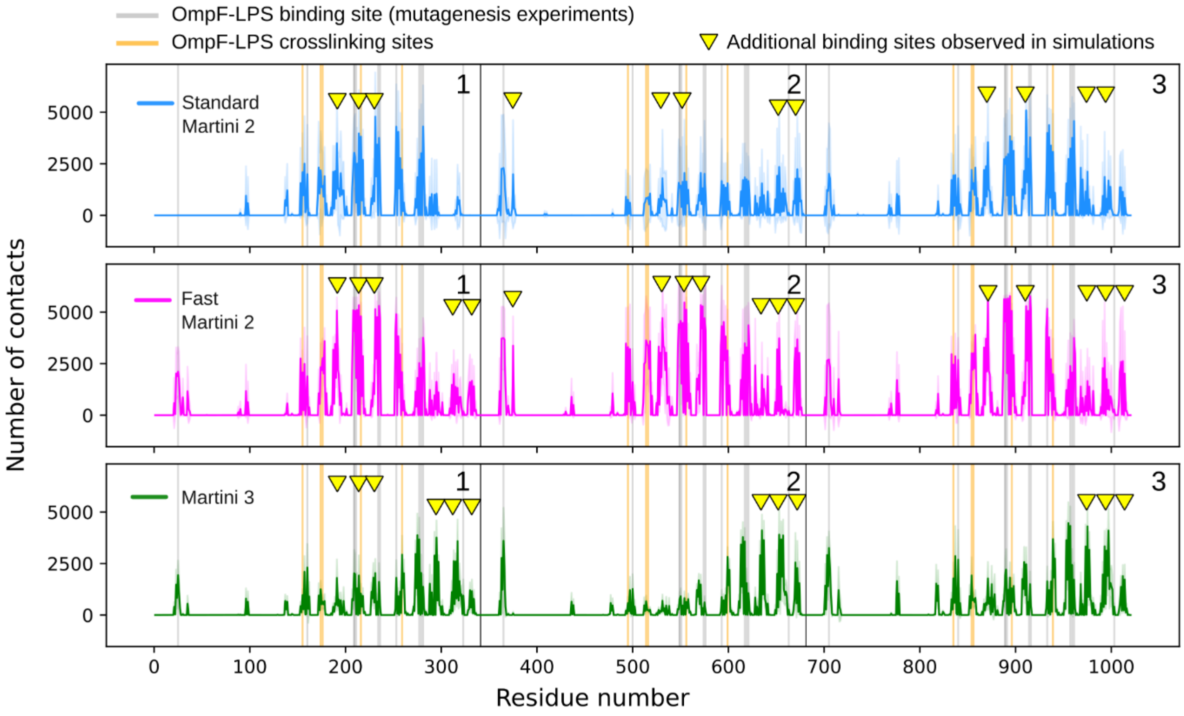

**S20. Number of OmpF-ReLPS contacts in standard Martini 2, fast Martini 2, and Martini 3 simulations with 2% ReLPS concentration in the outer leaflet.** For standard Martini 2 (upper panel), fast Martini 2 (middle), and Martini 3 (bottom), the contacts are shown in blue, magenta, and green respectively. Black lines are shown to differentiate between the residues of OmpF monomers. Grey lines indicate the residues that are proposed to contribute to LPS by mutagenesis experiments [14]. Orange lines indicate the residues that are involved in LPS binding as shown by OmpF-LPS crosslinking experiments [15]. The yellow triangles indicate newly identified OmpF residues from simulations that show a large number of contacts with ReLPS in addition to the residues that align with experimental data.

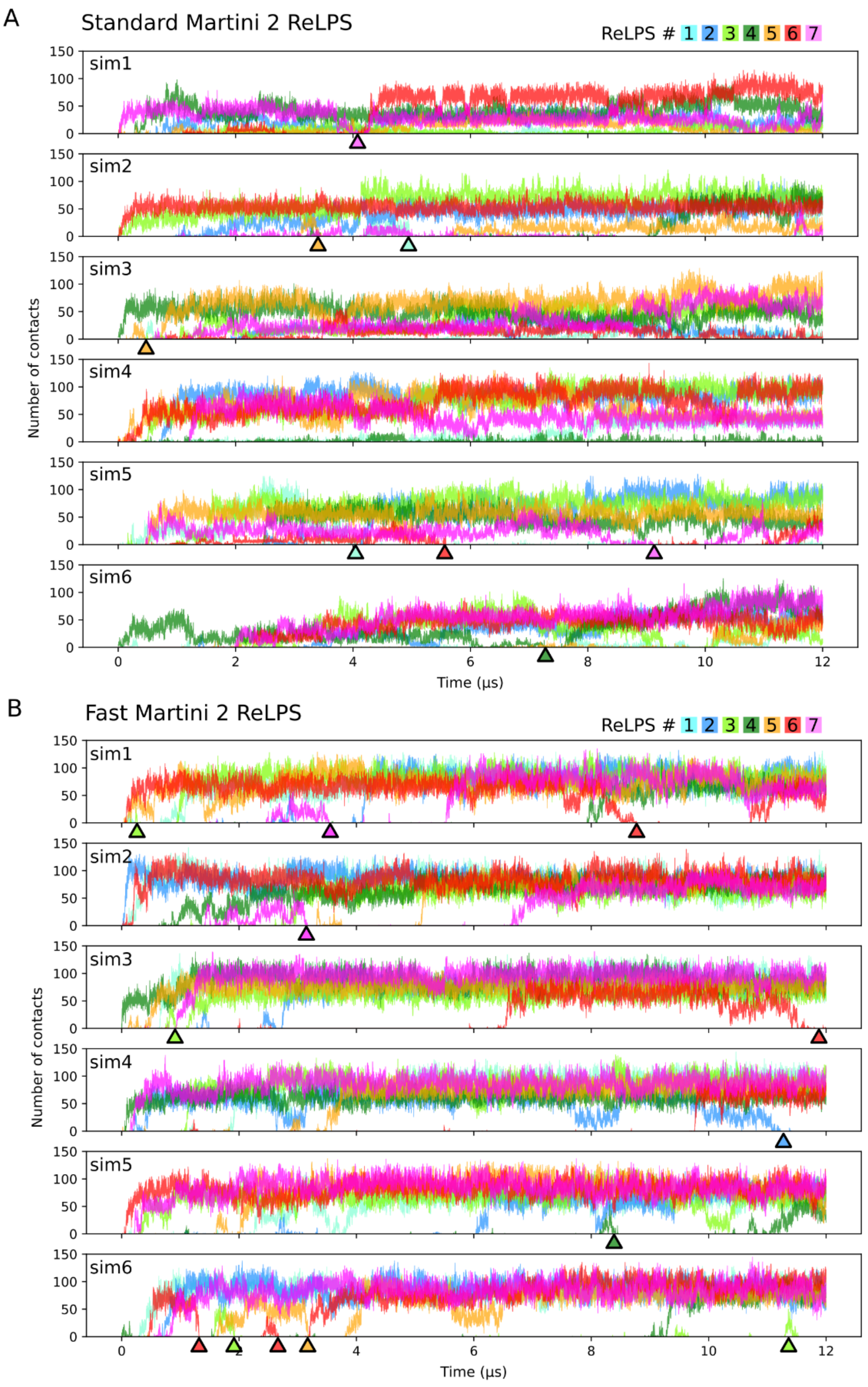

**S21. Dissociation events between Martini 2 ReLPS models and OmpF in an OM model with 2% ReLPS in the outer leaflet.** Number of OmpF-ReLPS contacts versus time shown from three individual simulations conducted with **(A)** standard and **(B)** fast Martini 2 ReLPS. Some dissociation events are indicated by arrows below the respective plots with their colours matching with that of the ReLPS molecule number.

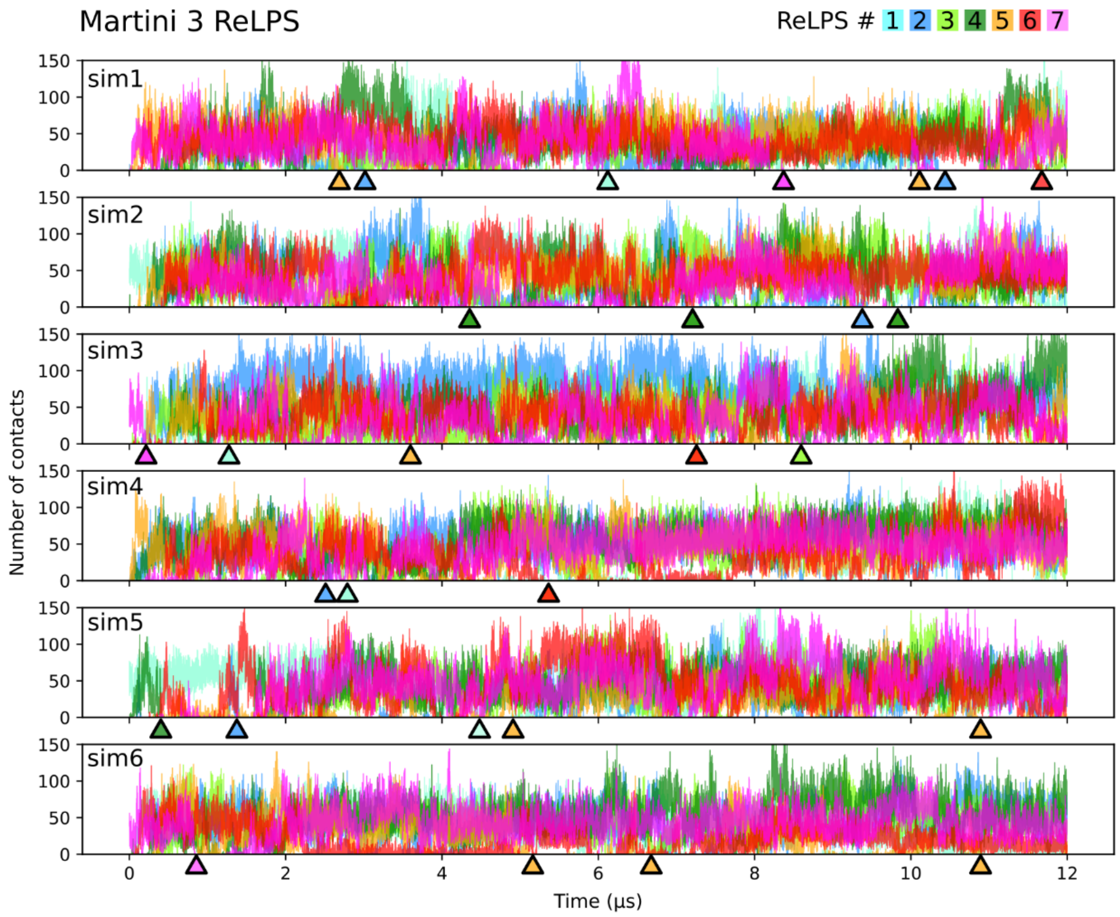

**S22. Dissociation events between our Martini 3 ReLPS model and OmpF in an OM model with 2% ReLPS in the outer leaflet**. Number of OmpF-ReLPS contacts versus time shown from three individual simulations conducted with our Martini 3 ReLPS model. Some dissociation events are indicated by arrows below the respective plots with their colours matching with that of the ReLPS molecule number.

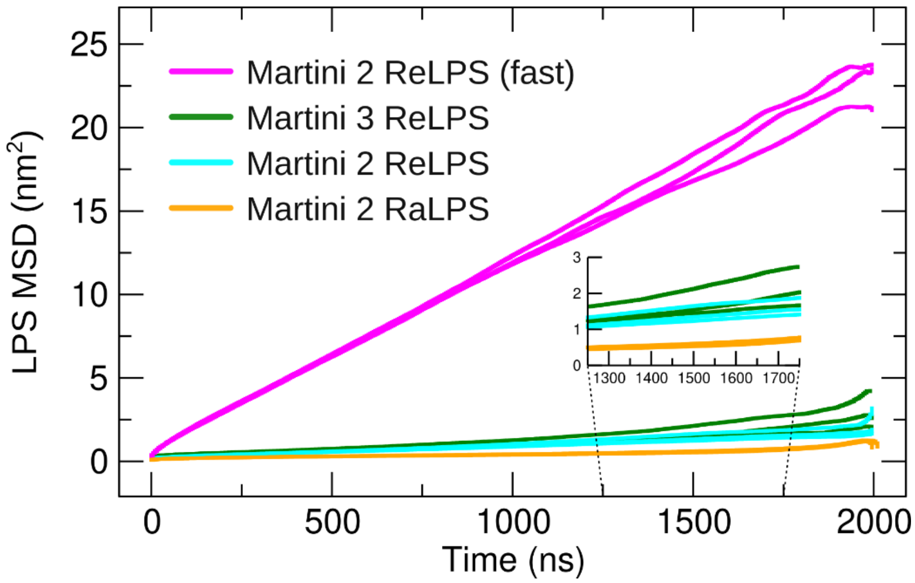

**S23. Comparing the lateral diffusion of the standard Martini 2 ReLPS and RaLPS, fast Martini 2 ReLPS, our Martini 3 ReLPS models.** This was done by computing their MSD versus time from the OMP island simulations. The lateral MSDs computed from each simulation replica are plotted individually.
