## additional text within the supplementary information for "Faster but not sweeter: A model of *E. coli* Re-level lipopolysaccharide for Martini 3 and a Martini 2 version with accelerated kinetics": README.md

# martini_3_ReLPS
These are the parameters for ReLPS to be used with the Martini 3 force field

Please cite the following paper when using this model: 
"Faster but not sweeter: A model of E. coli Re-level lipopolysaccharide for Martini 3 and a Martini 2 version with accelerated kinetics", A.F. Brandner & D. Prakaash, A. Blanco Gonzalez, F. Waterhouse, S. Khalid (in preparation).
